## Supplementary material for "Spatial and Texture Analysis of Root System Distribution with Earth Mover’s Distance (STARSEED)": Source file: bmc_article.bib

% bmc_article.bib
% 
%  An example of bibtex entries.
%  Entries taken from BMC instructions for authors page.

% uncomment next line to make author-year bibliography
% @settings{label, options="nameyear"}

@article{blank,
    author  = {}, 
    title   = {},
    journal = {}, 
    year    = {},
    month   = {}, 
    volume  = {}, 
    number  = {}, 
    pages   = {},
    note    = {} 
}    

% Article within a journal
@article{koon,
    author  = {Koonin, E V and Altschul, S F and P Bork}, 
    title   = {BRCA1 protein products: functional motifs}, 
    journal = {Nat. Genet.}, 
    year    = {1996},
    volume  = {13}, 
    pages   = {266-267}
}    

%%%%%%%%
% Article within conference proceedings
@inproceedings{xjon,
    author    = {X Jones}, 
    title     = {Zeolites and synthetic mechanisms},
    booktitle = {Proceedings of the First National Conference on 
                Porous Sieves: 27-30 June 1996; Baltimore},
    year      = {1996},
    editor    = {Y Smith}, 
    pages     = {16-27},
}    

%%%%%%%%
%  Book chapter, or article within a book
@incollection{schn,
    author    = {E Schnepf}, 
    title     = {From prey via endosymbiont to plastids: 
             comparative studies in dinoflagellates},
    booktitle = {Origins of Plastids}, 
    editor    = {R A Lewin}, 
    publisher = {Chapman and Hall},
    pages     = {53-76}, 
    year      = {1993},
    address = {New York}, 
    edition = {2nd} 
}    

%%%%%%%%
% Complete book
@book{marg,
    author    = {L Margulis}, 
    title     = {Origin of Eukaryotic Cells},
    publisher = {Yale University Press}, 
    year      = {1970},
    address   = {New Haven} 
}


%%%%%%%%
% PHD Thesis
@phdthesis{koha,
    author = {R Kohavi}, 
    title  = {Wrappers for performance enhancement and
             obvious decision graphs},
    school = {Stanford University, Computer Science Department},
    year   = {1995}
}

%%%%%%%%
%  Miscellaneous: webpage link/urL, etc/
@misc{issnic,
    author  = {{ISSN International Centre}},
    title = {The ISSN register},
    url = {http://www.issn.org},
    year = {2006},
    urldate={Accessed 20 Feb 2007}
}

%%%%%%%%%%%%%%%%Branching Paper$$$$$$$$$
@article{brummer2021branching,
  title={Branching principles of animal and plant networks identified by combining extensive data, machine learning and modelling},
  author={Brummer, Alexander B and Lymperopoulos, Panagiotis and Shen, Jocelyn and Tekin, Elif and Bentley, Lisa P and Buzzard, Vanessa and Gray, Andrew and Oliveras, Imma and Enquist, Brian J and Savage, Van M},
  journal={Journal of the Royal Society Interface},
  volume={18},
  number={174},
  pages={20200624},
  year={2021},
  publisher={The Royal Society}
}

%%%%%%%%%%%%%%%%%% Dimensionality Reduction %%%%%%%%%%%%%%%%
@InProceedings{Oza2001DecorrelationDimReduction,
author="Oza, Nikunj C.
and Tumer, Kagan",
editor="Kittler, Josef
and Roli, Fabio",
title="Input Decimation Ensembles: Decorrelation through Dimensionality Reduction",
booktitle="Multiple Classifier Systems",
year="2001",
publisher="Springer Berlin Heidelberg",
address="Berlin, Heidelberg",
pages="238--247",
isbn="978-3-540-48219-2"
}

@article{Becht2019Dimensionality,
  title={Dimensionality reduction for visualizing single-cell data using UMAP},
  author={Becht, Etienne and McInnes, Leland and Healy, John and Dutertre, Charles-Antoine and Kwok, Immanuel WH and Ng, Lai Guan and Ginhoux, Florent and Newell, Evan W},
  journal={Nature biotechnology},
  volume={37},
  number={1},
  pages={38--44},
  year={2019},
  publisher={Nature Publishing Group}
}
@article{VanDerMaaten2008Visualizing,
  title={Visualizing data using t-SNE},
  author={Maaten, Laurens van der and Hinton, Geoffrey},
  journal={Journal of machine learning research},
  volume={9},
  number={Nov},
  pages={2579--2605},
  year={2008}
}

%%%%%%%XAI%%%%%%%%%
@article{gunning2017explainable,
	title={Explainable artificial intelligence (xai)},
	author={Gunning, David},
	journal={Defense Advanced Research Projects Agency (DARPA), nd Web},
	year={2017}
}
@article{adadi2018peeking,
	title={Peeking inside the black-box: A survey on Explainable Artificial Intelligence (XAI)},
	author={Adadi, Amina and Berrada, Mohammed},
	journal={IEEE Access},
	volume={6},
	pages={52138--52160},
	year={2018},
	publisher={IEEE}
}

%%%%%%%Root Architecture Analysis%%%%%%%%%%
@article{xu2020overcoming,
  title={Overcoming small minirhizotron datasets using transfer learning},
  author={Xu, Weihuang and Yu, Guohao and Zare, Alina and Zurweller, Brendan and Rowland, Diane L and Reyes-Cabrera, Joel and Fritschi, Felix B and Matamala, Roser and Juenger, Thomas E},
  journal={Computers and Electronics in Agriculture},
  volume={175},
  pages={105466},
  year={2020},
  publisher={Elsevier}
}

@article{smith2020segmentation,
  title={Segmentation of roots in soil with U-Net},
  author={Smith, Abraham George and Petersen, Jens and Selvan, Raghavendra and Rasmussen, Camilla Ru{\o}},
  journal={Plant Methods},
  volume={16},
  number={1},
  pages={1--15},
  year={2020},
  publisher={BioMed Central}
}

@article{yasrab2019rootnav,
  title={RootNav 2.0: Deep learning for automatic navigation of complex plant root architectures},
  author={Yasrab, Robail and Atkinson, Jonathan A and Wells, Darren M and French, Andrew P and Pridmore, Tony P and Pound, Michael P},
  journal={GigaScience},
  volume={8},
  number={11},
  pages={giz123},
  year={2019},
  publisher={Oxford University Press}
}

@article{yasrab2020rootnet,
  title={RootNet: A Convolutional Neural Networks for Complex Plant Root Phenotyping from High-Definition Datasets},
  author={Yasrab, Robail and Pound, Michael and French, Andrew and Pridmore, Tony},
  journal={bioRxiv},
  year={2020},
  publisher={Cold Spring Harbor Laboratory}
}

@article{yu2020root,
  title={Root identification in minirhizotron imagery with multiple instance learning},
  author={Yu, Guohao and Zare, Alina and Sheng, Hudanyun and Matamala, Roser and Reyes-Cabrera, Joel and Fritschi, Felix B and Juenger, Thomas E},
  journal={Machine Vision and Applications},
  volume={31},
  number={6},
  pages={1--13},
  year={2020},
  publisher={Springer}
}

@inproceedings{yu2020weakly,
  title={Weakly Supervised Minirhizotron Image Segmentation with MIL-CAM},
  author={Yu, Guohao and Zare, Alina and Xu, Weihuang and Matamala, Roser and Reyes-Cabrera, Joel and Fritschi, Felix B and Juenger, Thomas E},
  booktitle={European Conference on Computer Vision},
  pages={433--449},
  year={2020},
  organization={Springer}
}

@article{chung2020image,
  title={Image-Based Machine Learning Characterizes Root Nodule in Soybean Exposed to Silicon},
  author={Chung, Yong Suk and Lee, Unseok and Heo, Seong and Silva, Renato Rodrigues and Na, Chae-In and Kim, Yoonha},
  journal={Frontiers in plant science},
  volume={11},
  year={2020},
  publisher={Frontiers Media SA}
}
@article{pound2017deep,
  title={Deep machine learning provides state-of-the-art performance in image-based plant phenotyping},
  author={Pound, Michael P and Atkinson, Jonathan A and Townsend, Alexandra J and Wilson, Michael H and Griffiths, Marcus and Jackson, Aaron S and Bulat, Adrian and Tzimiropoulos, Georgios and Wells, Darren M and Murchie, Erik H and others},
  journal={Gigascience},
  volume={6},
  number={10},
  pages={gix083},
  year={2017},
  publisher={Oxford University Press}
}
@article{zhi2017root,
  title={Root growth and spatial distribution characteristics for seedlings raised in substrate and transplanted cotton},
  author={Zhi, Xiaoyu and Han, Yingchun and Li, Yabing and Wang, Guoping and Feng, Lu and Yang, Beifang and Fan, Zhengyi and Lei, Yaping and Du, Wenli and Mao, Shuchun},
  journal={Plos one},
  volume={12},
  number={12},
  pages={e0190032},
  year={2017},
  publisher={Public Library of Science San Francisco, CA USA}
}

@article{LI201747,
title = {Spatial distribution of soil water, soil temperature, and plant roots in a drip-irrigated intercropping field with plastic mulch},
journal = {European Journal of Agronomy},
volume = {83},
pages = {47-56},
year = {2017},
issn = {1161-0301},
author = {Xianyue Li and Jiří Šimůnek and Haibin Shi and Jianwen Yan and Zunyuan Peng and Xuewen Gong},
publisher={Elsevier}
}

@article{li2017effects,
  title={Effects of root spatial distribution on the elastic-plastic properties of soil-root blocks},
  author={Li, Yunpeng and Wang, Yunqi and Wang, Yujie and Ma, Chao},
  journal={Scientific Reports},
  volume={7},
  number={1},
  pages={1--11},
  year={2017},
  publisher={Nature Publishing Group}
}

%%%%%%%%Texture%%%%%%%%%%%%%%%%%%%
@article{frigui2008detection,
	title={Detection and discrimination of land mines in ground-penetrating radar based on edge histogram descriptors and a possibilistic $ k $-nearest neighbor classifier},
	author={Frigui, Hichem and Gader, Paul},
	journal={IEEE Transactions on Fuzzy Systems},
	volume={17},
	number={1},
	pages={185--199},
	year={2008},
	publisher={IEEE}
}
@inproceedings{dalal2005histograms,
	title={Histograms of oriented gradients for human detection},
	author={Dalal, Navneet and Triggs, Bill},
	booktitle={international Conference on computer vision \& Pattern Recognition (CVPR'05)},
	volume={1},
	pages={886--893},
	year={2005},
	organization={IEEE Computer Society}
}

@ARTICLE{peeples2021histogram,
  author={Peeples, Joshua and Xu, Weihuang and Zare, Alina},
  journal={IEEE Transactions on Artificial Intelligence}, 
  title={Histogram Layers for Texture Analysis}, 
  year={2022},
  volume={3},
  number={4},
  pages={541-552},
  doi={10.1109/TAI.2021.3135804}}
  
@article{tuceryan1993texture,
  title={Texture analysis},
  author={Tuceryan, Mihran and Jain, Anil K},
  journal={Handbook of pattern recognition and computer vision},
  pages={235--276},
  year={1993},
  publisher={World Scientific}
}

@article{liu2019bow,
  title={From BoW to CNN: Two decades of texture representation for texture classification},
  author={Liu, Li and Chen, Jie and Fieguth, Paul and Zhao, Guoying and Chellappa, Rama and Pietik{\"a}inen, Matti},
  journal={International Journal of Computer Vision},
  volume={127},
  number={1},
  pages={74--109},
  year={2019},
  publisher={Springer}
}
@inproceedings{cula2001compact,
  title={Compact representation of bidirectional texture functions},
  author={Cula, Oana G and Dana, Kristin J},
  booktitle={Proceedings of the 2001 IEEE Computer Society Conference on Computer Vision and Pattern Recognition. CVPR 2001},
  volume={1},
  pages={I--I},
  year={2001},
  organization={IEEE}
}
%%%%%%%%Superpixels%%%%%%%%%%%%%%%%%%%
@article{achanta2012slic,
  title={SLIC superpixels compared to state-of-the-art superpixel methods},
  author={Achanta, Radhakrishna and Shaji, Appu and Smith, Kevin and Lucchi, Aurelien and Fua, Pascal and S{\"u}sstrunk, Sabine},
  journal={IEEE transactions on pattern analysis and machine intelligence},
  volume={34},
  number={11},
  pages={2274--2282},
  year={2012},
  publisher={IEEE}
}
%%%%%%%%Roots%%%%%%%%%%%%%%%%%
@article{falk2020computer,
  title={Computer vision and machine learning enabled soybean root phenotyping pipeline},
  author={Falk, Kevin G and Jubery, Talukder Z and Mirnezami, Seyed V and Parmley, Kyle A and Sarkar, Soumik and Singh, Arti and Ganapathysubramanian, Baskar and Singh, Asheesh K},
  journal={Plant methods},
  volume={16},
  number={1},
  pages={5},
  year={2020},
  publisher={Springer}
}
@article{jorgensen2014spatial,
  title={Spatial root distribution of plants growing in vertical media for use in living walls},
  author={J{\o}rgensen, Lars and Dresb{\o}ll, Dorte Bodin and Thorup-Kristensen, Kristian},
  journal={Plant and soil},
  volume={380},
  number={1},
  pages={231--248},
  year={2014},
  publisher={Springer}
}
@article{lynch2007roots,
  title={Roots of the second green revolution},
  author={Lynch, Jonathan P},
  journal={Australian Journal of Botany},
  volume={55},
  number={5},
  pages={493--512},
  year={2007},
  publisher={CSIRO}
}
@article{atkinson2019uncovering,
  title={Uncovering the hidden half of plants using new advances in root phenotyping},
  author={Atkinson, Jonathan A and Pound, Michael P and Bennett, Malcolm J and Wells, Darren M},
  journal={Current opinion in biotechnology},
  volume={55},
  pages={1--8},
  year={2019},
  publisher={Elsevier}
}
@article{wu2018optimizing,
  title={Optimizing soil-coring strategies to quantify root-length-density distribution in field-grown maize: virtual coring trials using 3-D root architecture models},
  author={Wu, Qian and Wu, Jie and Zheng, Bangyou and Guo, Yan},
  journal={Annals of botany},
  volume={121},
  number={5},
  pages={809--819},
  year={2018},
  publisher={Oxford University Press US}
}
@article{zhu2011lab,
  title={From lab to field, new approaches to phenotyping root system architecture},
  author={Zhu, Jinming and Ingram, Paul A and Benfey, Philip N and Elich, Tedd},
  journal={Current opinion in plant biology},
  volume={14},
  number={3},
  pages={310--317},
  year={2011},
  publisher={Elsevier}
}
%%%%%%%%Root Modelling%%%%%%%%%%%%%%%%%
@article{shahzad2018ez,
  title={EZ-Root-VIS: a software pipeline for the rapid analysis and visual reconstruction of root system architecture},
  author={Shahzad, Zaigham and Kellermeier, Fabian and Armstrong, Emily M and Rogers, Simon and Lobet, Guillaume and Amtmann, Anna and Hills, Adrian},
  journal={Plant physiology},
  volume={177},
  number={4},
  pages={1368--1381},
  year={2018},
  publisher={Am Soc Plant Biol}
}
@article{tron2015can,
  title={Can diversity in root architecture explain plant water use efficiency? A modeling study},
  author={Tron, Stefania and Bodner, Gernot and Laio, Francesco and Ridolfi, Luca and Leitner, Daniel},
  journal={Ecological modelling},
  volume={312},
  pages={200--210},
  year={2015},
  publisher={Elsevier}
}
@article{armengaud2009ez,
  title={EZ-Rhizo: integrated software for the fast and accurate measurement of root system architecture},
  author={Armengaud, Patrick and Zambaux, Kevin and Hills, Adrian and Sulpice, Ronan and Pattison, Richard J and Blatt, Michael R and Amtmann, Anna},
  journal={The Plant Journal},
  volume={57},
  number={5},
  pages={945--956},
  year={2009},
  publisher={Wiley Online Library}
}
@article{postma2017opensimroot,
  title={OpenSimRoot: widening the scope and application of root architectural models},
  author={Postma, Johannes A and Kuppe, Christian and Owen, Markus R and Mellor, Nathan and Griffiths, Marcus and Bennett, Malcolm J and Lynch, Jonathan P and Watt, Michelle},
  journal={New Phytologist},
  volume={215},
  number={3},
  pages={1274--1286},
  year={2017},
  publisher={Wiley Online Library}
}
@article{zhao2017root,
  title={Root architecture simulation improves the inference from seedling root phenotyping towards mature root systems},
  author={Zhao, Jiangsan and Bodner, Gernot and Rewald, Boris and Leitner, Daniel and Nagel, Kerstin A and Nakhforoosh, Alireza},
  journal={Journal of experimental botany},
  volume={68},
  number={5},
  pages={965--982},
  year={2017},
  publisher={Oxford University Press UK}
}
@article{pages2020metamodelling,
  title={Metamodelling a 3D architectural root-system model to provide a simple model based on key processes and species functional groups},
  author={Pag{\`e}s, Lo{\"\i}c and Pointurier, Olivia and Moreau, Delphine and Voisin, Anne-Sophie and Colbach, Nathalie},
  journal={Plant and Soil},
  pages={1--21},
  year={2020},
  publisher={Springer}
}
@article{landl2018measuring,
  title={Measuring root system traits of wheat in 2D images to parameterize 3D root architecture models},
  author={Landl, Magdalena and Schnepf, Andrea and Vanderborght, Jan and Bengough, A Glyn and Bauke, Sara L and Lobet, Guillaume and Bol, Roland and Vereecken, Harry},
  journal={Plant and soil},
  volume={425},
  number={1},
  pages={457--477},
  year={2018},
  publisher={Springer}
}
@article{schnepf2018statistical,
  title={Statistical characterization of the root system architecture model CRootBox},
  author={Schnepf, Andrea and Huber, Katrin and Landl, Magdalena and Meunier, Felicien and Petrich, Lukas and Schmidt, Volker},
  journal={Vadose zone journal},
  volume={17},
  number={1},
  pages={1--11},
  year={2018},
  publisher={Wiley Online Library}
}

@article{pierret2013ij_rhizo,
  title={IJ\_Rhizo: an open-source software to measure scanned images of root samples},
  author={Pierret, Alain and Gonkhamdee, Santimaitree and Jourdan, Christophe and Maeght, Jean-Luc},
  journal={Plant and soil},
  volume={373},
  number={1},
  pages={531--539},
  year={2013},
  publisher={Springer}
}

@article{lynch1997simroot,
  title={SimRoot: modelling and visualization of root systems},
  author={Lynch, Jonathan P and Nielsen, Kai L and Davis, Robert D and Jablokow, Andrei G},
  journal={Plant and Soil},
  volume={188},
  number={1},
  pages={139--151},
  year={1997},
  publisher={Springer}
}

@article{schnepf2018crootbox,
  title={CRootBox: a structural--functional modelling framework for root systems},
  author={Schnepf, Andrea and Leitner, Daniel and Landl, Magdalena and Lobet, Guillaume and Mai, Trung Hieu and Morandage, Shehan and Sheng, Cheng and Z{\"o}rner, Mirjam and Vanderborght, Jan and Vereecken, Harry},
  journal={Annals of botany},
  volume={121},
  number={5},
  pages={1033--1053},
  year={2018},
  publisher={Oxford University Press US}
}
%%%%%%%%Texture Analysis%%%%%%%%%%%%%%%%%%%%%%%

%%%%%%%%Fractal Dimension and Lacunarity%%%%%%%

@book{mandelbrot1982fractal,
  title={The fractal geometry of nature},
  author={Mandelbrot, Benoit B and Mandelbrot, Benoit B},
  volume={1},
  year={1982},
  publisher={WH freeman New York}
}

@article{mandelbrot1967long,
  title={How long is the coast of Britain? Statistical self-similarity and fractional dimension},
  author={Mandelbrot, Benoit},
  journal={science},
  volume={156},
  number={3775},
  pages={636--638},
  year={1967},
  publisher={American Association for the Advancement of Science}
}

@article{keller1989texture,
  title={Texture description and segmentation through fractal geometry},
  author={Keller, James M and Chen, Susan and Crownover, Richard M},
  journal={Computer Vision, Graphics, and image processing},
  volume={45},
  number={2},
  pages={150--166},
  year={1989},
  publisher={Elsevier}
}

@article{mandelbrot1968fractional,
  title={Fractional Brownian motions, fractional noises and applications},
  author={Mandelbrot, Benoit B and Van Ness, John W},
  journal={SIAM review},
  volume={10},
  number={4},
  pages={422--437},
  year={1968},
  publisher={SIAM}
}

@incollection{voss1991random,
  title={Random fractals: characterization and measurement},
  author={Voss, Richard F},
  booktitle={Scaling phenomena in disordered systems},
  pages={1--11},
  year={1991},
  publisher={Springer}
}

@article{sarkar1992efficient,
  title={An efficient approach to estimate fractal dimension of textural images},
  author={Sarkar, Nirupam and Chaudhuri, Bidyut Baran},
  journal={Pattern recognition},
  volume={25},
  number={9},
  pages={1035--1041},
  year={1992},
  publisher={Elsevier}
}

@inproceedings{feng1996fractional,
  title={Fractional box-counting approach to fractal dimension estimation},
  author={Feng, Jie and Lin, Wei-Chung and Chen, Chin-Tu},
  booktitle={Proceedings of 13th international conference on Pattern recognition},
  volume={2},
  pages={854--858},
  year={1996},
  organization={IEEE}
}

@article{plotnick1993lacunarity,
  title={Lacunarity indices as measures of landscape texture},
  author={Plotnick, Roy E and Gardner, Robert H and O'Neill, Robert V},
  journal={Landscape ecology},
  volume={8},
  number={3},
  pages={201--211},
  year={1993},
  publisher={Springer}
}

%%%%%%%Calinski-Harabasz%%%%%%%%%%%%
@article{calinski1974dendrite,
  title={A dendrite method for cluster analysis},
  author={Cali{\'n}ski, Tadeusz and Harabasz, Jerzy},
  journal={Communications in Statistics-theory and Methods},
  volume={3},
  number={1},
  pages={1--27},
  year={1974},
  publisher={Taylor \& Francis}
}

%%%%%%%Earth Mover's Distance%%%%%%%%%%%%
@inproceedings{rubner1997earth,
  title={The earth mover’s distance, multi-dimensional scaling, and color-based image retrieval},
  author={Rubner, Yossi and Guibas, Leonidas J and Tomasi, Carlo},
  booktitle={Proceedings of the ARPA image understanding workshop},
  volume={661},
  pages={668},
  year={1997}
}

@inproceedings{rubner1998metric,
  title={A metric for distributions with applications to image databases},
  author={Rubner, Yossi and Tomasi, Carlo and Guibas, Leonidas J},
  booktitle={Sixth International Conference on Computer Vision (IEEE Cat. No. 98CH36271)},
  pages={59--66},
  year={1998},
  organization={IEEE}
}

@inproceedings{gulrajani2017improved,
  title={Improved training of wasserstein GANs},
  author={Gulrajani, Ishaan and Ahmed, Faruk and Arjovsky, Martin and Dumoulin, Vincent and Courville, Aaron},
  booktitle={Proceedings of the 31st International Conference on Neural Information Processing Systems},
  pages={5769--5779},
  year={2017}
}

@article{hitchcock1941distribution,
  title={The distribution of a product from several sources to numerous localities},
  author={Hitchcock, Frank L},
  journal={Journal of mathematics and physics},
  volume={20},
  number={1-4},
  pages={224--230},
  year={1941},
  publisher={Wiley Online Library}
}

@online{fisher2004earth,
  author = {Bob Fisher},
  title = {The Earth Mover's Distance},
  year = 2004,
  url = {https://homepages.inf.ed.ac.uk/rbf/CVonline/LOCAL_COPIES/RUBNER/emd.htm}
}

@article{benamou2000computational,
  title={A computational fluid mechanics solution to the Monge-Kantorovich mass transfer problem},
  author={Benamou, Jean-David and Brenier, Yann},
  journal={Numerische Mathematik},
  volume={84},
  number={3},
  pages={375--393},
  year={2000},
  publisher={Springer}
}
@inproceedings{zhang2020deepemd,
  title={DeepEMD: Few-Shot Image Classification With Differentiable Earth Mover's Distance and Structured Classifiers},
  author={Zhang, Chi and Cai, Yujun and Lin, Guosheng and Shen, Chunhua},
  booktitle={Proceedings of the IEEE/CVF Conference on Computer Vision and Pattern Recognition},
  pages={12203--12213},
  year={2020}
}
@inproceedings{kusner2015word,
  title={From word embeddings to document distances},
  author={Kusner, Matt and Sun, Yu and Kolkin, Nicholas and Weinberger, Kilian},
  booktitle={International conference on machine learning},
  pages={957--966},
  year={2015},
  organization={PMLR}
}
@article{salti2014shot,
  title={SHOT: Unique signatures of histograms for surface and texture description},
  author={Salti, Samuele and Tombari, Federico and Di Stefano, Luigi},
  journal={Computer Vision and Image Understanding},
  volume={125},
  pages={251--264},
  year={2014},
  publisher={Elsevier}
}
%%%%%%%%%%%%%%MDS%%%%%%%%%%%%%%%%%%
@article{kruskal1964multidimensional,
  title={Multidimensional scaling by optimizing goodness of fit to a nonmetric hypothesis},
  author={Kruskal, Joseph B},
  journal={Psychometrika},
  volume={29},
  number={1},
  pages={1--27},
  year={1964},
  publisher={Springer}
}

%%%%%%%%%%%%%%Fractal Analysis of Roots%%%%%%%%%%%%%%%%%%
@article{tatsumi1989fractal,
  title={Fractal analysis of plant root systems},
  author={TATSUMI, JIRO and YAMAUCHI, AKIRA and KONO, YASUHIRO},
  journal={Annals of Botany},
  volume={64},
  number={5},
  pages={499--503},
  year={1989},
  publisher={Oxford University Press}
}

@article{wang2009fractal,
  title={Fractal analysis on root systems of rice plants in response to drought stress},
  author={Wang, Hong and Siopongco, Joel and Wade, Len J and Yamauchi, Akira},
  journal={Environmental and Experimental Botany},
  volume={65},
  number={2-3},
  pages={338--344},
  year={2009},
  publisher={Elsevier}
}
@article{li2020fractal,
  title={Fractal and Topological Analyses and Antioxidant Defense Systems of Alfalfa (Medicago sativa L.) Root System under Drought and Rehydration Regimes},
  author={Li, Shuo and Wan, Liqiang and Nie, Zhongnan and Li, Xianglin},
  journal={Agronomy},
  volume={10},
  number={6},
  pages={805},
  year={2020},
  publisher={Multidisciplinary Digital Publishing Institute}
}
@article{walk2004modelling,
  title={Modelling applicability of fractal analysis to efficiency of soil exploration by roots},
  author={Walk, Thomas C and Van Erp, Erik and Lynch, Jonathan P},
  journal={Annals of Botany},
  volume={94},
  number={1},
  pages={119--128},
  year={2004},
  publisher={Oxford University Press}
}
@article{wu2014integrated,
  title={An integrated method for quantifying root architecture of field-grown maize},
  author={Wu, Jie and Guo, Yan},
  journal={Annals of botany},
  volume={114},
  number={4},
  pages={841--851},
  year={2014},
  publisher={Oxford University Press}
}
@article{eshel1998fractal,
  title={On the fractal dimensions of a root system},
  author={Eshel, A},
  journal={Plant, Cell \& Environment},
  volume={21},
  number={2},
  pages={247--251},
  year={1998},
  publisher={Wiley Online Library}
}
%%%%%%%%Sesame%%%%%%%%%%%%%%%%%
@article{gloaguen2021plasticity,
  title={The plasticity of early root development in Sesamum indicum L. as influenced by genotype, water, and nutrient availability},
  author={Gloaguen, Romain M and Brym, Zachary T and Peeples, Joshua and Xu, Weihuang and Chun, Hyen-Chung and Rowland, Diane L},
  journal={Rhizosphere},
  pages={100457},
  year={2021},
  publisher={Elsevier}
}

@incollection{ISLAM2016135,
 title = {Chapter 6 - Sesame},
 editor = {Surinder Kumar Gupta},
 booktitle = {Breeding Oilseed Crops for Sustainable Production},
 publisher = {Academic Press},
 address = {San Diego},
 pages = {135-147},
 year = {2016},
 isbn = {978-0-12-801309-0},
 doi = {https://doi.org/10.1016/B978-0-12-801309-0.00006-9},
 url = {https://www.sciencedirect.com/science/article/pii/B9780128013090000069},
 author = {Faisal Islam and Rafaqat A. Gill and Basharat Ali and Muhammad A. Farooq and Ling Xu and Ullah Najeeb and Weijun Zhou},
}

@article{anee2019oxidative,
  title={Oxidative damage and antioxidant defense in Sesamum indicum after different waterlogging durations},
  author={Anee, Taufika Islam and Nahar, Kamrun and Rahman, Anisur and Mahmud, Jubayer Al and Bhuiyan, Tasnim Farha and Alam, Mazhar Ul and Fujita, Masayuki and Hasanuzzaman, Mirza},
  journal={Plants},
  volume={8},
  number={7},
  pages={196},
  year={2019},
  publisher={Multidisciplinary Digital Publishing Institute}
}

@article{mensah2006simulated,
  title={Simulated flooding and drought effects on germination, growth, and yield parameters of sesame (Sesamum indicum L.)},
  author={Mensah, JK and Obadoni, BO and Eruotor, PG and Onome-Irieguna, F},
  journal={African Journal of Biotechnology},
  volume={5},
  number={13},
  year={2006}
}

@article{hamedani2020variability,
  title={Variability of root traits in sesame genotypes under different irrigation regimes},
  author={Hamedani, Nima Ghasemi and Gholamhoseini, Majid and Bazrafshan, Forrod and Amiri, Bahram and Habibzadeh, Farhad},
  journal={Rhizosphere},
  volume={13},
  pages={100190},
  year={2020},
  publisher={Elsevier}
}

@article{bahrami2012effect,
  title={Effect of drought stress on germination and seedling growth of sesame cultivars (Sesamum indicum L.).},
  author={Bahrami, Helale and Razmjoo, Jamshid and Jafari, A Ostadi and others},
  journal={International Journal of AgriScience},
  volume={2},
  number={5},
  pages={423--428},
  year={2012},
  publisher={International Academic Journals}
}

@article{dargahi2013effect,
  title={Effect of water deficit stress on root morphological characters in sesame cultivars},
  author={Dargahi, Y and Asghari, A and Shokrpour, M and Rasoulzadeh, A},
  year={2013},
  publisher={ELECTRONIC JOURNAL OF CROP PRODUCTION}
}

@article{sarkar2016effect,
  title={Effect of duration of water-logging on crop stand and yield of sesame},
  author={Sarkar, PK and Khatun, A and Singha, A},
  journal={International Journal of Innovation and Applied Studies},
  volume={14},
  number={1},
  pages={1},
  year={2016},
  publisher={Citeseer}
}

@article{wei2013morpho,
  title={Morpho-anatomical and physiological responses to waterlogging of sesame (Sesamum indicum L.)},
  author={Wei, Wenliang and Li, Donghua and Wang, Linhai and Ding, Xia and Zhang, Yanxin and Gao, Yuan and Zhang, Xiurong},
  journal={Plant science},
  volume={208},
  pages={102--111},
  year={2013},
  publisher={Elsevier}
}

@article{manschadi2006role,
  title={The role of root architectural traits in adaptation of wheat to water-limited environments},
  author={Manschadi, Ahmad M and Christopher, John and deVoil, Peter and Hammer, Graeme L},
  journal={Functional plant biology},
  volume={33},
  number={9},
  pages={823--837},
  year={2006},
  publisher={CSIRO}
}


%%%%%%%%Statistics%%%%%%%%%%%%%%%%%
@article{LenthR2020,
  title={Emmeans: Estimated marginal means, aka least-squares means.},
  author={Lenth, R.},
  url = {https://CRAN.R-project.org/package=emmeans},
}

@article{hothorn2008simultaneous,
  title={Simultaneous inference in general parametric models},
  author={Hothorn, Torsten and Bretz, Frank and Westfall, Peter},
  journal={Biometrical Journal: Journal of Mathematical Methods in Biosciences},
  volume={50},
  number={3},
  pages={346--363},
  year={2008},
  publisher={Wiley Online Library}
}

%%%%%Tools for root tracing%%%%%%%%%%%%%
@article{ingram2001software,
  title={Software for measuring root characters from digital images},
  author={Ingram, Keith T and Leers, Gary A},
  journal={Agronomy Journal},
  volume={93},
  number={4},
  pages={918--922},
  year={2001},
  publisher={Wiley Online Library}
}

@article{abramoff2004image,
  title={Image processing with ImageJ},
  author={Abr{\`a}moff, Michael D and Magalh{\~a}es, Paulo J and Ram, Sunanda J},
  journal={Biophotonics international},
  volume={11},
  number={7},
  pages={36--42},
  year={2004},
  publisher={Laurin Publishing}
}

@article{le2010dart,
  title={DART: a software to analyse root system architecture and development from captured images},
  author={Le Bot, Jacques and Serra, Val{\'e}rie and Fabre, Jos{\'e} and Draye, Xavier and Adamowicz, St{\'e}phane and Pag{\`e}s, Lo{\"\i}c},
  journal={Plant and Soil},
  volume={326},
  number={1},
  pages={261--273},
  year={2010},
  publisher={Springer}
}

@article{lobet2013online,
  title={An online database for plant image analysis software tools},
  author={Lobet, Guillaume and Draye, Xavier and P{\'e}rilleux, Claire},
  journal={Plant methods},
  volume={9},
  number={1},
  pages={1--8},
  year={2013},
  publisher={Springer}
}

@article{cai2015rootgraph,
  title={RootGraph: a graphic optimization tool for automated image analysis of plant roots},
  author={Cai, Jinhai and Zeng, Zhanghui and Connor, Jason N and Huang, Chun Yuan and Melino, Vanessa and Kumar, Pankaj and Miklavcic, Stanley J},
  journal={Journal of experimental botany},
  volume={66},
  number={21},
  pages={6551--6562},
  year={2015},
  publisher={Oxford University Press UK}
}
