## Supplementary material for "Spatial and Texture Analysis of Root System Distribution with Earth Mover’s Distance (STARSEED)": Source file: STARSEED_Plant_Methods_V2.tex

%% BioMed_Central_Tex_Template_v1.06
%%                                      %
%  bmc_article.tex            ver: 1.06 %
%                                       %

%%IMPORTANT: do not delete the first line of this template
%%It must be present to enable the BMC Submission system to
%%recognise this template!!

%%%%%%%%%%%%%%%%%%%%%%%%%%%%%%%%%%%%%%%%%
%%                                     %%
%%  LaTeX template for BioMed Central  %%
%%     journal article submissions     %%
%%                                     %%
%%          <8 June 2012>              %%
%%                                     %%
%%                                     %%
%%%%%%%%%%%%%%%%%%%%%%%%%%%%%%%%%%%%%%%%%

%%%%%%%%%%%%%%%%%%%%%%%%%%%%%%%%%%%%%%%%%%%%%%%%%%%%%%%%%%%%%%%%%%%%%
%%                                                                 %%
%% For instructions on how to fill out this Tex template           %%
%% document please refer to Readme.html and the instructions for   %%
%% authors page on the biomed central website                      %%
%% https://www.biomedcentral.com/getpublished                      %%
%%                                                                 %%
%% Please do not use \input{...} to include other tex files.       %%
%% Submit your LaTeX manuscript as one .tex document.              %%
%%                                                                 %%
%% All additional figures and files should be attached             %%
%% separately and not embedded in the \TeX\ document itself.       %%
%%                                                                 %%
%% BioMed Central currently use the MikTex distribution of         %%
%% TeX for Windows) of TeX and LaTeX.  This is available from      %%
%% https://miktex.org/                                             %%
%%                                                                 %%
%%%%%%%%%%%%%%%%%%%%%%%%%%%%%%%%%%%%%%%%%%%%%%%%%%%%%%%%%%%%%%%%%%%%%

%%% additional documentclass options:
%  [doublespacing]
%  [linenumbers]   - put the line numbers on margins

%%% loading packages, author definitions

%\documentclass[twocolumn]{bmcart}% uncomment this for twocolumn layout and comment line below
\documentclass[double]{bmcart}

%%% Load packages
\usepackage{amsthm,amsmath}
%\RequirePackage[numbers]{natbib}
%\RequirePackage[authoryear]{natbib}% uncomment this for author-year bibliography
%\RequirePackage{hyperref}
\usepackage[utf8]{inputenc} %unicode support
%\usepackage[applemac]{inputenc} %applemac support if unicode package fails
%\usepackage[latin1]{inputenc} %UNIX support if unicode package fails
\usepackage{siunitx}
\usepackage{subcaption}
\usepackage{multirow}
\usepackage{url}
\usepackage{algorithm}
\usepackage{algpseudocode}
\usepackage{varwidth}
\usepackage{adjustbox,lipsum}
\renewcommand{\figurename}{Fig.}
\usepackage{bm}
\usepackage{soul}
\usepackage{color}
\usepackage{amsfonts}
\usepackage[switch]{lineno}
%\linenumbers

%%%%%%%%%%%%%%%%%%%%%%%%%%%%%%%%%%%%%%%%%%%%%%%%%
%%                                             %%
%%  If you wish to display your graphics for   %%
%%  your own use using includegraphic or       %%
%%  includegraphics, then comment out the      %%
%%  following two lines of code.               %%
%%  NB: These line *must* be included when     %%
%%  submitting to BMC.                         %%
%%  All figure files must be submitted as      %%
%%  separate graphics through the BMC          %%
%%  submission process, not included in the    %%
%%  submitted article.                         %%
%%                                             %%
%%%%%%%%%%%%%%%%%%%%%%%%%%%%%%%%%%%%%%%%%%%%%%%%%

% \def\includegraphic{}
% \def\includegraphics{}

%%% Put your definitions there:
\startlocaldefs
\endlocaldefs

%%% Begin ...
\begin{document}

%%% Start of article front matter
\begin{frontmatter}

\begin{fmbox}
\dochead{Methodology}

%%%%%%%%%%%%%%%%%%%%%%%%%%%%%%%%%%%%%%%%%%%%%%
%%                                          %%
%% Enter the title of your article here     %%
%%                                          %%
%%%%%%%%%%%%%%%%%%%%%%%%%%%%%%%%%%%%%%%%%%%%%%

\title{Spatial and Texture Analysis of Root System Distribution with Earth Mover's Distance (STARSEED)}

%%%%%%%%%%%%%%%%%%%%%%%%%%%%%%%%%%%%%%%%%%%%%%
%%                                          %%
%% Enter the authors here                   %%
%%                                          %%
%% Specify information, if available,       %%
%% in the form:                             %%
%%   <key>={<id1>,<id2>}                    %%
%%   <key>=                                 %%
%% Comment or delete the keys which are     %%
%% not used. Repeat \author command as much %%
%% as required.                             %%
%%                                          %%
%%%%%%%%%%%%%%%%%%%%%%%%%%%%%%%%%%%%%%%%%%%%%%

% \author[
%   addressref={aff1},                   % id's of addresses, e.g. {aff1,aff2}
%   corref={aff1},                       % id of corresponding address, if any
% % noteref={n1},                        % id's of article notes, if any
% ]{\inits{J.E.}\fnm{Jane E.} \snm{Doe}}
% \author[
%   addressref={aff1,aff2},
% ]{\inits{J.R.S.}\fnm{John R.S.} \snm{Smith}}

\author[
  addressref={aff1},                   % id's of addresses, e.g. {aff1,aff2}
  corref={aff1},                       % id of corresponding address, if any
% noteref={n1},                        % id's of article notes, if any
]{\inits{J.}\fnm{Joshua} \snm{Peeples}}
\author[
  addressref={aff2},
]{\inits{W.}\fnm{Weihuang} \snm{Xu}}
\author[
  addressref={aff3},
]{\inits{R.}\fnm{Romain} \snm{Gloaguen}}
\author[
  addressref={aff4},
]{\inits{D.}\fnm{Diane} \snm{Rowland}}
\author[
  addressref={aff2},
]{\inits{A.}\fnm{Alina} \snm{Zare}}
\author[
  addressref={aff6,aff5},
]{\inits{Z.}\fnm{Zachary} \snm{Brym}}

\address[id=aff1]{%
  \orgdiv{Department of Electrical and Computer Engineering},
  \orgname{Texas A\&M University},
%   \street{},
  \postcode{77845}
  \city{College Station},
  \cny{USA}
}
\address[id=aff2]{%
  \orgdiv{Department of Electrical and Computer Engineering},
  \orgname{University of Florida},
%   \street{},
  \postcode{32611}
  \city{Gainesville},
  \cny{USA}
}
\address[id=aff3]{%
  \orgdiv{InTerACT Research Unit},
  \orgname{UniLaSalle Bauvais},
%   \street{},
%   \postcode{32611}
%   \city{Gainesville},
  \cny{France}
}
\address[id=aff4]{%
  \orgdiv{College of Natural Sciences, Forestry, and Agriculture},
  \orgname{University of Maine},
%   \street{},
  \postcode{04469}
  \city{Orono},
  \cny{USA}
}
\address[id=aff6]{%
  \orgdiv{Tropical Research and Education Center},
  \orgname{University of Florida},
%   \street{},
  \postcode{33031}
  \city{Gainesville},
  \cny{USA}
}
\address[id=aff5]{%
  \orgdiv{Department of Agronomy},
  \orgname{University of Florida},
%   \street{},
  \postcode{32611}
  \city{Gainesville},
  \cny{USA}
}
%%%%%%%%%%%%%%%%%%%%%%%%%%%%%%%%%%%%%%%%%%%%%%
%%                                          %%
%% Enter short notes here                   %%
%%                                          %%
%% Short notes will be after addresses      %%
%% on first page.                           %%
%%                                          %%
%%%%%%%%%%%%%%%%%%%%%%%%%%%%%%%%%%%%%%%%%%%%%%

%\begin{artnotes}
%%\note{Sample of title note}     % note to the article
%\note[id=n1]{Equal contributor} % note, connected to author
%\end{artnotes}

\end{fmbox}% comment this for two column layout

%%%%%%%%%%%%%%%%%%%%%%%%%%%%%%%%%%%%%%%%%%%%%%%
%%                                           %%
%% The Abstract begins here                  %%
%%                                           %%
%% Please refer to the Instructions for      %%
%% authors on https://www.biomedcentral.com/ %%
%% and include the section headings          %%
%% accordingly for your article type.        %%
%%                                           %%
%%%%%%%%%%%%%%%%%%%%%%%%%%%%%%%%%%%%%%%%%%%%%%%

\begin{abstractbox}

\begin{abstract} % abstract
\textbf{Purpose:} Root system architectures are complex and challenging to characterize effectively for agronomic and ecological discovery.\newline 
\textbf{Methods:} We propose a new method, Spatial and Texture Analysis of Root SystEm distribution with Earth mover's Distance (STARSEED), for comparing root system distributions that incorporates spatial information through a novel application of the Earth Mover's Distance (EMD). \newline
\textbf{Results:} We illustrate that the approach captures the response of sesame root systems for different genotypes and soil moisture levels. STARSEED provides quantitative and visual insights into changes that occur in root architectures across experimental treatments.\newline
\textbf{Conclusion:} STARSEED can be generalized to other plants and provides insight into root system architecture development and response to varying growth conditions not captured by existing root architecture metrics and models. The code and data for our experiments are publicly available: \url{https://github.com/GatorSense/STARSEED}.

% \parttitle{First part title} %if any
% Text for this section.

% \parttitle{Second part title} %if any
% Text for this section.
\end{abstract}

%%%%%%%%%%%%%%%%%%%%%%%%%%%%%%%%%%%%%%%%%%%%%%
%%                                          %%
%% The keywords begin here                  %%
%%                                          %%
%% Put each keyword in separate \kwd{}.     %%
%%                                          %%
%%%%%%%%%%%%%%%%%%%%%%%%%%%%%%%%%%%%%%%%%%%%%%

\begin{keyword}
Root architecture, Earth Mover's Distance, Image Analysis, Sesamum indicum, Artificial Intelligence
\end{keyword}

% MSC classifications codes, if any
%\begin{keyword}[class=AMS]
%\kwd[Primary ]{}
%\kwd{}
%\kwd[; secondary ]{}
%\end{keyword}

\end{abstractbox}
%
%\end{fmbox}% uncomment this for two column layout

\end{frontmatter}

%%%%%%%%%%%%%%%%%%%%%%%%%%%%%%%%%%%%%%%%%%%%%%%%
%%                                            %%
%% The Main Body begins here                  %%
%%                                            %%
%% Please refer to the instructions for       %%
%% authors on:                                %%
%% https://www.biomedcentral.com/getpublished %%
%% and include the section headings           %%
%% accordingly for your article type.         %%
%%                                            %%
%% See the Results and Discussion section     %%
%% for details on how to create sub-sections  %%
%%                                            %%
%% use \cite{...} to cite references          %%
%%  \cite{koon} and                           %%
%%  \cite{oreg,khar,zvai,xjon,schn,pond}      %%
%%                                            %%
%%%%%%%%%%%%%%%%%%%%%%%%%%%%%%%%%%%%%%%%%%%%%%%%

%%%%%%%%%%%%%%%%%%%%%%%%% start of article main body
% <put your article body there>

%%%%%%%%%%%%%%%%
%% Background %%
%%
\section*{Introduction}
\label{intro}
Studying plant roots is one of the keys to achieving the second Green Revolution \cite{lynch2007roots}. The study of plant roots requires, among other things, effective characterization and development of comparative methods for root development, architecture, and spatial distribution, including early in the root system's development. Current methods, such as the widely used WinRHIZO\textsuperscript{TM} and WinRHIZO\textsuperscript{TM} Tron software suite (Regent Instruments Inc., Quebec, Canada) use 2D images of roots to measure individual parameters related to morphology and topology. These standard software packages have also been developed to further make accessible the analysis of individual root traits \cite{pierret2013ij_rhizo,armengaud2009ez}. These tools capture information relevant to characterize root spatial distribution (RSD). However, standard software packages typically provide parameters that are only capturing one or a few specific aspects of a root system such as total root length, surface area, branching angle and root order. These packages are less effective at holistically quantifying intact root systems and how they change in response to edaphic conditions. {To address this shortcoming, machine learning can be used in tandem with these biological features to automate and improve understanding of root systems \mbox{\cite{brummer2021branching}}}.

A subarea of machine learning, deep learning, has been widely used for automating root detection and segmentation from 2D images of root systems \cite{chung2020image,pound2017deep,yasrab2019rootnav,yu2020root,xu2020overcoming}. Deep learning methods can require high computational power and large amounts of annotated data however, as well as lack explainability \cite{gunning2017explainable,adadi2018peeking}. Yet these techniques have shown promise for localizing and identifying unique 2D texture features useful for root phenotyping \cite{pound2017deep}. Texture, meaning the spatial arrangement of the pixel values in a raster grid \cite{peeples2021histogram}, is a powerful cue that can be used to identify patterns tied to RSD characterization. A straightforward approach to compute texture from an image is to use counts (\textit{i.e.}, histograms) of local pixel intensities \cite{cula2001compact}. {This representation of texture can be defined as counting the percentage of root pixels in an area of the image. By counting the number of root pixels in each cell of a raster grid, this allows for quantification of localized pixel intensity depending on the cell size of the grid.} In addition to this baseline approach, one can compute more complex texture features such as fractal dimension and lacunarity. 

Root systems can be described as approximate fractal objects over a finite range of scale. A fractal object is defined by two fundamental characteristics: self-similarity (\textit{i.e.} shape variations of the object on one scale are repeated at another) \cite{mandelbrot1967long} and the possession of a fractal dimension that is expressed as a non-integer dimension, as opposed to the more familiar integer Euclidean dimensions \cite{mandelbrot1982fractal}. This fractal dimension value can be calculated for root systems using a specific approach called the box-counting method \cite{tatsumi1989fractal, eshel1998fractal} and has been used successfully to capture root system architecture \cite{tatsumi1989fractal,li2020fractal,wang2009fractal}. 
 
A related feature to fractal dimension is lacunarity. Lacunarity captures the distribution and size of ``gaps" in an object \mbox{\cite{mandelbrot1982fractal,plotnick1993lacunarity}}. An object, in our case the image of a root system, will be highly \textit{lacunar} if it contains many gaps distributed over a greater range. Lacunarity will thus be smaller in value when root systems are dense and increase with the presence of gaps {(\textit{i.e.}, areas of the image where roots are not located or areas of the growth media matrix where roots are not located)} and a coarse spatial arrangement of roots  \mbox{\cite{keller1989texture,mandelbrot1982fractal}}. Since some visually distinct images can have the same fractal dimension, lacunarity provides a feature that can aid in discriminating between these distinct textures \cite{keller1989texture,voss1991random,mandelbrot1968fractional}. 

The aforementioned texture features are promising candidates in building a new tool to accurately and precisely describe RSD. Therefore, a novel algorithm is needed that would allow for the calculation of these texture features from 2D root images and spatially explicit comparisons between root systems based on the feature values. For these aims, Earth Mover's Distance (EMD) appears to be a promising candidate approach \mbox{\cite{rubner1998metric}.} EMD, also known as the Wasserstein-1 distance \cite{gulrajani2017improved}, is used to determine quantitative differences between two distributions \cite{rubner1998metric}. EMD has several advantages such as allowing for partial matching (\textit{i.e.,} comparisons can be made between representations of different sizes such as comparing smaller and larger root systems) and matches our human visual perception when the chosen ground distances (\textit{i.e.}, distance between feature vectors) is meaningful \cite{rubner1998metric}. Essentially, we want to use EMD to find the minimal distance or amount of ``work" to transform one root spatial distribution to another. 

In this study, we developed the Spatial and Texture Analysis of Root SystEm distribution with Earth mover's Distance (STARSEED) approach, to better characterize RSD and overall root soil exploration. Our approach used EMD to quantify the heterogeneity among root structures. The proposed STARSEED approach allowed us to establish meaningful biological connections between the treatments such as cultivar and moisture level, and the observed RSD as well as provide detailed qualitative and quantitative analysis. The aims and contributions of STARSEED are the following: 1) characterize the spatial arrangements of roots in local (\textit{i.e.}, smaller regions of the image) and global (\textit{i.e.}, whole image) contexts, 2) extract useful information to describe the distribution of roots in the image, and 3) give precise insight for biological interpretation.

\section*{Materials and Methods}
\label{sec:1}
\subsection*{\textbf{Greenhouse Setup and Data Collection}}
Data presented and used in this work is coming from a previous study that has been published \mbox{\cite{gloaguen2021plasticity}}. A summary of the greenhouse setup and data collection of the materials and methods are presented here. Sesame was grown in 64 custom-made rhizoboxes. Each rhizobox was filled with $1550$g of inert calcine clay (Turface Athletics, Buffalo Grove, IL, USA), henceforth referred to as soil. A single seed from one of four non-dehiscent sesame cultivars, all provided by Sesaco Inc. (Sesaco32, Sesaco35, Sesaco38 and Sesaco40, referred to as S32, S35, S38 and S40 thereafter), was planted per box. Four soil water content treatments were implemented: $60$, $80$, $100$ and $120\%$ of the soil water holding capacity, corresponding to $688$, $918$, $1147$ and $1376$ mL of water per rhizobox, respectively. The soil and water were mixed thoroughly together before filling the rhizoboxes to promote homogeneity of the soil water content throughout. The top of each rhizobox was covered with Press’N Seal Cling Wrap (Glad, Oakland, CA, USA) to minimize water evaporation. No water was added afterwards throughout the experiment with the assumption that soil water content, in the absence of evaporation, stayed relatively constant throughout the 16-21 days of each run experiment. A small hole was pierced in the plastic wrap upon seedling emergence to allow for leaf and stem growth. The two factors were arranged in a complete randomized design with 4 replications in a greenhouse where the daily temperature was maintained between 25-35 degrees Celsius.  

Four runs of 64 rhizoboxes were completed. Run 1 and 2 were prepared with soil and water only, while run 3 and 4 were fertilized with $1.51mg$ of 15-5-15 + Ca + Mg Peters Excel mix fertilizer (ICL Specialty Fertilizers, Summerville SC) and $0.29g$ of ammonium sulfate were dissolved in the water applied to each rhizobox prior to mixing with the soil. Baking soda was added as needed to neutralize the fertilizer solution to a pH of 6. Plants were grown for a duration of 21 days after planting (DAP) for the runs without fertilizer and 16 DAP for the runs with added fertilizer due to their fast growth.  

On the last day of the experiment, all rhizoboxes were scanned with a Plustek OpticSlim 1180 A3 flatbed scanner (Plustek Inc., Santa Fe, CA, USA) to generate the raw images with the size $6800\times4676$ at a resolution of 400 Dots Per Inch (DPI). Rhizoboxes were all set on the scanner the same way using a physical guide so that all rhizoboxes had the same position in the scanned images. For each run, images from seeds that did not germinate or seedlings that died during the experiment were not considered in the analysis; 11 images were thus removed from Run 1, 10 from Run 2, 11 from Run 3 and 5 from Run 4. The total number of analyzed images was 107 Runs 1 and 2, and 113 for Run 3 and 4. {Representative RGB image samples for each cultivar and water level are shown in supplemental Figures \mbox{\ref{fig:S3}} and \mbox{\ref{fig:S4}}.} The roots from all analyzed images were hand-traced with WinRHIZOTron\textsuperscript{TM} (Regent Instruments Inc., Quebec, Canada) and total root length (TRL) was obtained.

\subsection*{\textbf{Spatial and Texture Analysis of Root SystEm distribution with Earth mover's Distance (STARSEED)}}
The overall pipeline of our proposed method is illustrated in Figure \mbox{\ref{fig:method}} and explained in more detail in Algorithm \mbox{\ref{alg:STARSEED}}. {STARSEED consisted of five major steps. The first step was to take the input images and perform preprocessing to isolate the roots pixels from the background to focus on the architecture. For step 2, we divided the image into equal-sized bins by generating local regions of the image to improve computational efficiency and incorporate spatial information. Step 3 involved extracting texture features to describe the root pixels within each bin and generate a compact representation of the image (\textit{i.e.}, signature). The fourth step used EMD to measure the magnitude and direction of changes between each root signature. Lastly, we projected the pairwise distance matrix to perform qualitative and quantitative analysis of the root images and their corresponding treatment.} Details and rationale behind each step of this algorithm are provided in the following subsections.

 \begin{figure*}[htb]
    \centering
    \includegraphics[width=0.95\linewidth]{Figures/Figure1.png}
    \caption{Overview of the proposed STARSEED method step by step.}
    \label{fig:method}
\end{figure*}

\begin{algorithm}[htb]
	\caption{STARSEED Overall Process}
	\label{alg:STARSEED}
	\begin{algorithmic}[1]
	\Require $N$ input images {$\bm{X} \in \mathbb{R}^{ H \times W}$ , $N$ corresponding treatment labels $y \in \mathbb{R}^{1}$, $B$ number of spatial bins, $g_{\bm{\theta}}(\cdot)$ preprocessing function, $f_{\bm{\theta}}(\cdot)$ texture feature extraction, $d$ embedding dimension}
	\State \begin{varwidth}[t]{\linewidth}
	    \hskip\algorithmicindent{Preprocess images, $\bm{X}_{processed} \gets g_
	    {\bm{\theta}}(\bm{X})$} \
    \State \hskip\algorithmicindent Partition each image into $B$ regions \
    \State \hskip\algorithmicindent Compute feature and centroid location, $\bm{X}_{features} \gets f_{\bm{\theta}}(\bm{X}_{processed})$ for each region to get a  signature, $\bm{P} \in \mathbb{R}^{B\times 3}$, for each image\
    \State \hskip\algorithmicindent Compute EMD score and flow matrix between each image signature using Equation \ref{eq:EMD} to generate pairwise 
    distance matrix, $\bm{D} \in \mathbb{R}^{N \times N}$ \
    \State \hskip\algorithmicindent Project $\bm{D}$ using MDS \cite{kruskal1964multidimensional} into d-dimensional feature space to compute embedding, $\bm{E} \in \mathbb{R}^{N \times d} $
    \State \hskip\algorithmicindent Compute Calinski-Harabasz (CH) score using $\bm{E}$ and $y$ %$\bm{Y}$ 
	\end{varwidth}
	\State {\Return EMD Scores, CH Index, Flow Matrices}
	\end{algorithmic}
\end{algorithm}


\paragraph*{\textbf{Image Preprocessing - Step 1}} The initial step of the proposed approach was to perform preprocessing of the images to isolate the root pixels from the background. Raw images were manually traced using WinRHIZO\textsuperscript{TM} Tron software to generate the labels for root pixels in each image. The black and white binary images were reconstructed based on these labels, in which pixels that corresponded to roots were assigned a value of \textit{1} in the mask and the non-root pixels were assigned a value of \textit{0}. The binary images were then cropped and calibrated, and downsampled by a factor of eight through average pooling and smoothed by a Gaussian filter ($\sigma=1$) to mitigate noise before extracting texture features.

\paragraph*{\textbf{Sub-region Generation and Feature/Signature Extraction - Steps 2 \& 3}}

In order to apply EMD to characterize and compare RSDs using STARSEED, we constructed a signature representation, a composition of texture features computed spatially, for each image. Since texture is undefined at a single point \cite{tuceryan1993texture}, a local neighborhood needed to be identified to compute the texture features comprising the signature. In our approach, we divided each image into bins of equal sizes using a grid, each bin serving as the local neighborhood. Within each bin, texture features were computed to locally characterize the RSD. The size of the grid/bins provides trade-offs between computational efficiency and localization. For example, a larger grid yielding large bins corresponds to less computation cost, but each texture feature is computed over a larger area, resulting in a loss of localization. {The computational time for the method scaled linearly with the number of bins used (e.g., more bins corresponded to increased computational time) and this information is captured in Supplemental Figures \mbox{\ref{fig:S1}} and \mbox{\ref{fig:S2}}. For the proposed method, the grid size that corresponded to the maximum CH score was used to further analyze STARSEED as discussed in the following Experiments and Results section.}

\begin{figure*}[htb]
    \centering
    \includegraphics[width=0.95\linewidth]{Figures/Figure2.jpg}

    \caption{For step 2, the image is divided into $B$ equally-sized bins after preprocessing. Next, the feature/signature extraction for step 3 is completed by representing each bin as a $\mathbb{R}^{3}$ vector that consist of the bin center location ($x$ and $y$) and the texture feature ($f$) describing the pixels in the bin. The set of these $B$ vectors is the final signature representation. The signature can also be represented as a heatmap that shows the feature value for the pixels in a region. In this example, the percentage of root pixels is used as the texture feature.}
    \label{fig:Step_2}
\end{figure*}

Each bin of the image was represented through a cluster representative. The cluster representative consisted of spatial coordinates (\textit{i.e.}, horizontal and vertical position of the bin center) and a weight to create a $\mathbb{R}^3$ vector to represent a local spatial bin of the image. The weight given to each bin was computed by the texture feature extracted from the pixels in the bin. {An example of steps 2 and 3 is shown in Figure {\ref{fig:Step_2}}.} We calculated three feature values in each bin: percentage of root pixels, fractal dimension, and lacunarity. To calculate the percentage of root pixels, we computed the total number of root pixels present in each bin divided by the area (\textit{i.e.}, total number of pixels) of that same bin. The percentage of root pixels  provides direct insight into the distribution of roots in the image. If a region of the image has denser roots, the percentage of root pixels in the bins present in that region will be higher. The final representation of the image was a signature, which was the set of clusters from the image. By constructing our signature in this manner, we incorporated spatial (locally at the bin level and globally over the whole image) and texture information, to represent the root distributions.

\paragraph*{\textbf{Novel Earth Mover's Distance Application - Step 4}}
EMD can be used to effectively evaluate certain characteristics between different distributions, including images \cite{rubner1998metric}. Images are comprised of pixels and these pixels can be clustered or assigned to meaningful groups based on shared characteristics such as spatial location. The set of these clusters are used to form a \textit{signature}, a more compact representation of the image to increase computational efficiency \cite{rubner1998metric}.
Given an image with $C$ clusters, the signature representation, $P$, is given by Equation \ref{eq:signature} where $\mathbf{p}_i$ is the cluster representative and $w_{\mathbf{p}_i}$ is the weight of the cluster: 
\begin{equation}
    P = \{(\mathbf{p}_1,w_{\mathbf{p}_1}),...,(\mathbf{p}_C,w_{\mathbf{p}_C})\}. \label{eq:signature}
\end{equation}
Typically, the cluster representative is a feature vector and the weight is the percentage of pixels in a cluster expressing that feature \cite{rubner1998metric}. While the selection of the cluster representative is application dependent, defining the cluster representative as the information/descriptor (\textit{i.e.}, describing features) and importance (\textit{i.e.}, weight) provides a clear interpretation for EMD within disparate areas of an image. In our proposed STARSEED method, the signature representations were generated as mentioned in Step 3.

Once the signature representations are constructed, we used EMD to generate a pairwise distance matrix $\bm{D}$ between each pair of RSD signatures, choosing one RSD signature as the reference within each pair. After determining the magnitude of changes between two clusters, the  minimal flow matrix, $\mathbf{F}$, is used to compute EMD \cite{rubner1998metric}. Given two image signatures, $P$ and $Q$, with $S$ and $T$ representatives respectively, EMD is formulated in Equation \ref{eq:EMD} where $d_{ij}$ is the ground distance between the centroid of regions $\mathbf{p}_i$ and $\mathbf{q}_j$ and $f_{ij}$ is the optimal flow between $\mathbf{p}_i$ and $\mathbf{q}_j$:
\begin{equation}
    EMD(P,Q) = \frac{\sum_{i=1}^{S}\sum_{j=1}^{T}d_{ij}f_{ij}}{\sum_{i=1}^{S}\sum_{j=1}^{T}f_{ij}} \label{eq:EMD}
\end{equation}

EMD captures the dissimilarity between spatial distributions: larger values indicate more dissimilarity or ``work" to move the defined ``earth" feature to the cluster representative. EMD allows one to measure the global change between two spatial distributions (\textit{i.e.}, distance measure) as well as local changes between the two sources of information through the flow matrix, $\bm{F}$. If we can translate a 2D root image into a spatial distribution of the values for a given texture feature calculated from this image, EMD would then be able to measure the changes in this distribution from one root system image to the next. Consequently, EMD would provide a two-level comparison between these distributions: one at the global scale (\textit{i.e.}, the whole distribution or the whole image) that we can term ``holistic" through the EMD measure, and one at the local scale through the flow matrix that shows the distinct magnitude and direction of changes between each image. 

In our experiments, we first computed EMD between each pair of images, separately between Runs 1-2 and Runs 3-4. Regions containing no roots were removed from our calculations to mitigate the impact of background on our results. Therefore, we were comparing root signatures of different sizes. In order to not favor smaller signatures, the normalization factor was added to the EMD calculation as shown in Equation \ref{eq:EMD}. Any distance can be used to define the ground distance, $d_{ij}$ \cite{rubner1998metric}. To consider the root spatial distribution, we compared the center location of each spatial bin by selecting the Euclidean distance as our ground distance to compute $d_{ij}$.

\begin{figure*}[htb!]
    \centering
    \includegraphics[width=0.95\linewidth]{Figures/Figure3.png}

    \caption{Example of qualitative results produced by projecting EMD matrix through MDS for the percentage of root pixels as the feature and 2000 spatial bins for images from Runs 3 and 4. MDS served as a visualization approach to show the relationships in the pairwise distance matrix, $D$. The RSD images are arranged in the 2D space such that images with similar RSD are near one another (\textit{i.e.}, similar MDS coordinates).{Additional analysis can be completed by incorporating the moisture or cultivar information. The CH index (step 6) is computed to assess the MDS projection. The different colored frames represent the different moisture levels. The CH indices for 60\%, 80\%, 100\%, and 120\% are 8.67, 9.59, 0.23, and 55.68 respectively. The overall CH index for this projection is 15.57. The 120\% water level is the most different from the other architectures and the 120\% water level is the most compact and separated class visually as well as quantitatively through the CH index.}}
    \label{fig:MDS_roots}
\end{figure*}

\paragraph*{\textbf{Projection Using MDS - Step 5}} Once the distances between each image signature representation were computed to generate the pairwise distance matrix, we can use a dimensionality reduction approach to project the matrix into two dimensions. We projected the data into two dimensions to qualitatively and quantitatively evaluate the pairwise distance matrix. We used multi-dimensional scaling (MDS) \cite{kruskal1964multidimensional} as this method had been shown to work well with EMD to identify patterns among images that share some characteristics \cite{rubner1998metric,rubner1997earth}.
% \begin{figure*}[htb!]
%     \centering
%     \includegraphics[width=0.95\linewidth]{Figures/Figure3.png}

%     \caption{Example of qualitative results produced by projecting EMD matrix through MDS for the percentage of root pixels as the feature and 2000 spatial bins for images from Runs 3 and 4. MDS served as a visualization approach to show the relationships in the pairwise distance matrix, $D$. The RSD images are arranged in the 2D space such that images with similar RSD are near one another (\textit{i.e.}, similar MDS coordinates). }
%     \label{fig:MDS_roots}
% \end{figure*}
 To illustrate step 5 in the algorithm, Figure \ref{fig:MDS_roots} provides a visualization of the EMD matrix projection in a 2D space through MDS for Run 3 and Run 4 with the percentage of root pixels as the feature calculated for the highest number of spatial bins (2000 in each image). The output from the MDS method is an $\mathbb{R} ^{N \times 2}$ embedding matrix, $E$, that is a projection of the pairwise distance matrix in the Euclidean space. Once $E$ is computed, the embedding is then used in Step 6 to calculate Calinski-Harabasz (CH) index \cite{calinski1974dendrite} to quantitatively assess relationships among the root images when considering the treatment values (cultivar and water levels). 


\paragraph*{\textbf{Assessment of Relationships for RSD - Steps 6}} 
Lastly, we calculated the CH \cite{calinski1974dendrite} index to measure the intra-cluster similarity (\textit{i.e.}, RSD representatives of the same cultivar or water level) and inter-cluster dissimilarity (\textit{i.e.}, RSD representatives of different cultivar or water level) across treatment levels in each pair of runs. By using the EMD approach, we were able to score the differences between each treatment level, highlighting: a) the magnitude and b) the location of the RSD dissimilarities.
Ideally, samples that belong to the same cluster should be ``close" (\textit{i.e.}, smaller EMD distances or intra-cluster variance) while samples from different clusters should be ``well-separated" (\textit{i.e.}, larger EMD distances or inter-cluster variance). When the clusters are dense and well-separated, the CH index is larger. Finally, a non-parametric Kruskal-Wallis test was performed on the CH scores within each treatment for each pair of runs across features, and then separately for each feature across cultivars or across water levels. When the test was rejected ($p<.05$), Dunnett's test with Bonferroni's adjustment was used to compare treatment levels with one another.

\section*{Experiments and Results}
\label{sec:Results}
Our dataset consisted of 220 images across four different runs. {To select the best bin size of for the experiments, we investigated the impact of coarser (\textit{i.e.}, small number of spatial bins - large grid) and finer (\textit{i.e.}, large number of spatial bins - small grid) scales. The number of spatial bins per image varied from $100$ to $2000$ in steps of $100$ for a total of 20 values. Then, we performed quantitative and qualitative analysis for local and global RSD supported by the STARSEED method.} For our local analysis, we calculated the CH index using different features to evaluate the effectiveness of these extracted features at capturing RSD differences between different growing conditions and cultivars. For our global analysis, we aggregated each centered root image to create a representative for each treatment to elucidate the RSD spatial differences between these representative images. We discuss the results of the local and global analysis of STARSEED in the {following} sections. 


\subsection*{\textbf{Local Analysis: Relationships to Treatment Level Representatives}} \label{sec:local}

The average scores and standard deviations of the CH indices across all number of spatial bins for Runs 1 and 2 are shown in Table \ref{tab:CH_run12} while the scores for Runs 3 and 4 are shown in Table \ref{tab:CH_run34}. The small standard deviations indicate that our method was robust across the sizes of spatial bins (\textit{i.e.}, the scale) used for our analysis. The statistical analysis of the CH scores showed that \% root pixels tended to be the feature with the most discriminating power as evidenced by the highest CH score out of the three features considered, except for both fractal dimension and lacunarity on cultivar in Runs 1 and 2. For the rest of this study, we displayed the EMD results for the \% root pixels feature, except when presenting results for the cultivar differences in Runs 1 and 2 where we will use the fractal dimension feature. {The number of spatial bins that corresponded to the highest CH score is shown in Table 3}.   
In Runs 1 and 2, the CH indices for water levels ($1.50$ overall average) were not as high ($p<.05$) as for cultivars ($3.96$ overall average), indicating more overall variability of RSD without added nutrients between cultivars than between moisture levels (Table \ref{tab:CH_run12}). When looking specifically at each treatment level's score, both the S32 cultivar and the 80\% water level had the largest CH indices within cultivar and water level treatments respectively, indicating these treatments had the most distinct RSD. For Runs 3 and 4 with added nutrients, the trends were inverse in that the average CH indices were higher across all three features and spatial bins for water level ($14.70$ overall average) than for cultivar ($1.62$ overall average). The highest average CH score was observed for the 120\% soil water level, indicating a distinct RSD for this treatment compared to the three other soil water treatments (Table \ref{tab:CH_run34}).

\begin{table*}[htb]
% table caption is above the table
\renewcommand{\arraystretch}{1.5}
\caption{Average CH Indices and their associated standard deviation for each feature by cultivar and water level for Runs 1 and 2 across all sizes of spatial bins. Error values are reported with $\pm1$ standard deviation. {Bold uppercase letters indicate differences in CH score between levels for either cultivar or water Level across all three texture features ($p<.05$). Bold lowercase letters indicate differences in CH score between the three features across all treatment levels for cultivar and water Level separately ($p<.05$).}}
%}
\label{tab:CH_run12}       % Give a unique label
% For LaTeX tables use
\begin{center}
%\resizebox{\columnwidth}{!}{%
\begin{adjustbox}{max width=\textwidth}
\begin{tabular}{|c|c|c|c|c|c|c|c|c|c|c|}
    \hline
    Feature & \multicolumn{5}{c|}{Cultivar} & \multicolumn{5}{c|}{Water Level} \\
    \hline
     & S32 \textbf{A} & S35 \textbf{C} & S38 \textbf{D} & S40 \textbf{B} & All & 60\% \textbf{C} & 80\% \textbf{A}& 100\% \textbf{B}& 120\% \textbf{B} & All\\
    \hline
    \hline
    \% Root Pixels & 11.07$\pm$0.43 & 0.64$\pm$0.08 & 0.62$\pm$0.07 & 2.79$\pm$0.16 & 3.41$\pm$0.08 \textbf{b} & 1.08$\pm$0.16 & 3.70$\pm$0.27 & 0.95$\pm$0.17 & 0.65$\pm$0.15 & 1.88$\pm$0.08  \textbf{a}\\
    \hline
    Fractal & 11.76$\pm$1.57 & 1.09$\pm$0.41 & 0.38$\pm$0.25 & 3.18$\pm$0.80 & 3.74$\pm$0.47 \textbf{a} & 0.21$\pm$0.21 & 3.02$\pm$0.38 & 0.86$\pm$0.39 & 1.64$\pm$0.61 & 1.57$\pm$0.17 \textbf{b}\\
    \hline
    Lacunarity & 11.37$\pm$1.58 & 1.04$\pm$0.35 & 0.31$\pm$0.23 & 3.30$\pm$0.82 & 3.67$\pm$0.47 \textbf{a} & 0.22$\pm$0.19 & 3.23$\pm$0.47 & 0.77$\pm$0.37 & 1.74$\pm$0.62 & 1.65$\pm$0.21 \textbf{b}\\
    \hline
\end{tabular}%
%}
\end{adjustbox}
\end{center}
\end{table*}

\begin{table*}[htb]
% table caption is above the table
\renewcommand{\arraystretch}{1.5}
\caption{Average CH Indices and their associated standard deviation for each feature by cultivar and water level for Runs 3 and 4 across all sizes of spatial bins. Error values are reported with $\pm1$ standard deviation. {Bold uppercase letters indicate differences in CH score between levels for either cultivar or water Level across all three texture features ($p<.05$). Bold lowercase letters indicate differences in CH score between the three features across all treatment levels for cultivar and water Level separately ($p<.05$).}}
\label{tab:CH_run34}       % Give a unique label
% For LaTeX tables use
\begin{center}
%\resizebox{\columnwidth}{!}{
\begin{adjustbox}{max width=\textwidth}
\begin{tabular}{|c|c|c|c|c|c|c|c|c|c|c|}
    \hline
    Feature & \multicolumn{5}{c|}{Cultivar} & \multicolumn{5}{c|}{Water Level} \\
    \hline
    {} & S32 \textbf{B} & S35 \textbf{A} & S38 \textbf{C} & S40 \textbf{A}
 & All & 60\% \textbf{B} & 80\% \textbf{B} & 100\% \textbf{C} & 120\% \textbf{A} & All\\
    \hline
    \hline
    \% Root Pixels & 1.46$\pm$0.07 & 2.57$\pm$0.16 & 1.40$\pm$0.03 & 1.98$\pm$0.11 & 1.79$\pm$0.06 \textbf{a} & 8.62$\pm$0.15 & 9.66$\pm$0.20 & 0.19$\pm$0.04 & 54.72$\pm$0.58 & 15.40$\pm$0.09 \textbf{a}\\
    \hline
    Fractal & 1.71$\pm$0.42 & 1.79$\pm$0.94 & 0.76$\pm$0.25 & 1.93$\pm$0.35 & 1.48$\pm$0.29 \textbf{b} & 6.00$\pm$0.67 & 6.31$\pm$1.15 & 2.40$\pm$1.03 & 38.13$\pm$7.58 & 12.70$\pm$1.78 \textbf{b}\\
    \hline
    Lacunarity & 1.61$\pm$0.49 & 1.77$\pm$0.8 & 0.66$\pm$0.29 & 1.80$\pm$0.36 & 1.38$\pm$0.31 \textbf{b} & 5.72$\pm$1.05 & 5.82$\pm$1.37 & 2.39$\pm$1.37 & 36.40$\pm$8.62 & 12.07$\pm$2.24 \textbf{b}\\
    \hline
\end{tabular}
\end{adjustbox}
%}
\end{center}
\end{table*}


\begin{table*}[htb]
% table caption is above the table
%\renewcommand{\arraystretch}{.5}
\caption{Number of spatial bins that scored the maximum CH index for each feature, cultivar, water level in each pair of runs.}
\label{tab:numSP}       % Give a unique label
% For LaTeX tables use
\begin{center}
%\resizebox{\columnwidth}{!}{%
\begin{tabular}{|c|c|c|c|c|}
    \hline
    Features & \multicolumn{2}{c|}{Run 1 and 2} & \multicolumn{2}{c|}{Run 3 and 4} \\
    \hline
    {} & Cultivar & Water Level & Cultivar & Water Level \\
    \hline
    \hline
    \%Root Pixels & 100 & 900 & 200 & 2000 \\
    \hline
    Fractal & 700 & 700 & 900 & 200\\
    \hline
    Lacunarity & 700 & 1000 & 1300 & 600\\
    \hline
\end{tabular}
%}
\end{center}
\end{table*}


\subsection*{\textbf{Global Analysis: EMD Between Treatments}}
\label{sec:global}
To identify global trends in RSD for each treatment, we aggregated each centered root image to create a representative for each treatment. For each representative root image, we computed the feature representation as shown in Figure \ref{fig:method} from the average feature values within each spatial bin for images with the same cultivar or water level. After we obtained these representatives, the EMD was computed between each cultivar or water level. By using the EMD approach, we highlighted the differences between each treatment level by displaying a) the magnitude and b) the location of the RSD dissimilarities, as shown in Figures \ref{fig:Run1S32} through \ref{fig:Run3_120}.

\begin{figure*}[htb]
    \centering
    \includegraphics[width=0.95\linewidth]{Figures/Figure4.png}
    \caption{RSD, EMD score, and value changes between S32 (Figure \ref{fig:Run1S32}a) and S35, S38 and S40 (Figure \ref{fig:Run1S32}b-d respectively) for Run 1 and 2. The color scale for Figure \ref{fig:Run1S32}a-d represents the fractal dimension value of each spatial bin. Figure \ref{fig:Run1S32}e shows the overall EMD score. Figure \ref{fig:Run1S32}f, Figure \ref{fig:Run1S32}g, and Figure \ref{fig:Run1S32}h represent the 20\% largest value changes between S32 and the three other cultivars; red spatial bins are unique to S32, green spatial bins are unique to S35, S38 or S40, and yellow spatial bins are common to both cultivars.}
    \label{fig:Run1S32}
\end{figure*}

The EMD calculation is able to capture the variability of the RSDs between cultivar and soil water levels within each pairs of runs, as illustrated by Figure \ref{fig:Run1S32}. The white arrows (Figures \ref{fig:Run1S32}-\ref{fig:Run3_120}f, g, h) represent the 20\% largest value changes between the reference treatment level (Figures \ref{fig:Run1S32}-\ref{fig:Run3_120}a) and the other levels (Figures \ref{fig:Run1S32}-\ref{fig:Run3_120}b, c, d). The EMD scores between the reference treatment level and the other levels are shown in Figures \ref{fig:Run1S32}-\ref{fig:Run3_120}e. Smaller EMD scores indicate higher similarity between the two RSDs. 

\paragraph*{\textbf{Differences between cultivars}}
Figure \ref{fig:Run1S32} shows the processed RSD images for each cultivar and three EMD results when these cultivars are compared with S32 as the reference cultivar. The EMD score (Figure \ref{fig:Run1S32}e) was smallest between S32 and S35, indicating that the two RSDs were most similar. This is confirmed by comparing Figure \ref{fig:Run1S32}f, Figure \ref{fig:Run1S32}g and Figure \ref{fig:Run1S32}h: Figure \ref{fig:Run1S32}f comparing S32 to S35 has the shortest and fewest arrows. Arrow abundance and length is a direct representation of the magnitude of the value changes between the two RSD. Lengthier and/or more numerous arrows are therefore indicative of more dissimilar RSDs compared with EMD results showing fewer and/or smaller arrows. Figure \ref{fig:Run1S32} shows that the changes in RSDs between S32 and S35 are relatively small, and for the most part are concentrated in the deeper section of the root system. Specifically, S35 seemed to have a more laterally spread out root distribution towards the deepest portion of the image, and a more concentrated root mass along the tap root right above that. Out of the three other cultivars, S32 was most different from S38, and Figure \ref{fig:Run1S32}g shows that the majority of the 20\% highest value changes in RSD between these cultivars were recorded toward the middle region of the image. S38 also appears to spread out more laterally in the mid-section of the image compared to S32. 

\begin{figure*}[htb]
    \centering
    \includegraphics[width=0.95\linewidth]{Figures/Figure5.png}
    \caption{RSD, EMD score and largest value changes between S40 (Figure \ref{fig:Run1S40}a) and S32, S35 and S38 (Figure \ref{fig:Run1S40}b-d respectively) for Run 1 and 2. The color scale for Figure \ref{fig:Run1S40}b-d represents the fractal dimension value of each spatial bin. Figure \ref{fig:Run1S40}e shows the EMD score overall. Figure \ref{fig:Run1S40}f, Figure \ref{fig:Run1S40}g, and Figure \ref{fig:Run1S40}h represent the top 20\% value changes between S40 and the three other cultivars; red spatial bins are unique to S40, green spatial bins are unique to S32, S35 or S38, and yellow spatial bins are common to both cultivars.}
    \label{fig:Run1S40}
\end{figure*}

The differences between S32 and S40 were located in the upper part of the root system, as shown by the location of the top 20\% value change between the two RSD in Figure \ref{fig:Run1S32}h. S32 tended to have a more spread out root distribution towards the top of the soil surface compared to S40. According to Table \ref{tab:CH_run12}, the RSD for S40 was different from all three other cultivars' RSD. The EMD score (Figure \ref{fig:Run1S40}) yet showed that the S40 RSD was further from the S32 RSD than the S35 and S38 RSD (Figure \ref{fig:Run1S40}e).

\begin{figure*}[htb]
    \centering
    \includegraphics[width=0.95\linewidth]{Figures/Figure8.png}
    \caption{RSD, EMD score and largest value changes between S40 (Figure \ref{fig:Run3S40}a) and S32, S35 and S38 (Figure \ref{fig:Run3S40}b-d respectively) for Run 3 and 4. The color scale for Figure \ref{fig:Run3S40}b-d represents the percentage of root pixels in each spatial bin. Figure \ref{fig:Run3S40}e shows the EMD score overall. Figure \ref{fig:Run3S40}f, Figure \ref{fig:Run3S40}g, and Figure \ref{fig:Run3S40}h represent the top 20\% value changes between S40 and the three other cultivars; red spatial bins are unique to S40, green spatial bins are unique to S32, S35 or S38, and yellow spatial bins are common to both cultivars.}
    \label{fig:Run3S40}
\end{figure*}
All cultivars appear to expand laterally with the addition of nutrients as shown in Figure \ref{fig:Run3S40}. S40 and S35 were the cultivars with the most distinct RSD as indicated by the highest CH score in Table \ref{tab:CH_run34}. Figure \ref{fig:Run3S40} shows specifically that for S40, the differences between this RSD and the three others remained mostly marginal: the EMD score remained low ($<$ 3.5 , Figure \ref{fig:Run3S40}e) and the 20\% largest value changes between the S40 RSD and the others only represented a few number of changes (Fig. \ref{fig:Run3S40}f, g, h). Interestingly, the RSD for S40 was most dissimilar to the RSD for S32 both with or without added nutrients. 


\paragraph*{\textbf{Differences between moisture levels}}
For Runs 1 and 2, when considering soil water levels, the RSD were overall very similar as shown in Table \ref{tab:CH_run12}. The RSD that was most distinct from the others was that of the 80\% moisture treatment (Figure \ref{fig:Run1_80}). The 80\% moisture treatment was overall most different from 100\%, and was the most similar to 120\% and the 60\% treatment RSD (Figure \ref{fig:Run1_80}e). Specifically, the 80\% treatment had a visibly denser central region than the three other moisture levels (Figure \ref{fig:Run1_80}a through Figure \ref{fig:Run1_80}d). This is reflected more clearly in Figure \ref{fig:Run1_80}f through h: the 80\% treatment  showed a more concentrated root distribution closer to the top of the root system compared to the other treatments, and so the arrows tend to point toward the upper center of the root system. 
\begin{figure*}[t]
    \centering
    \includegraphics[width=0.95\linewidth]{Figures/Figure7.png}
    \caption{RSD, EMD score and largest value changes between 80\% (Figure \ref{fig:Run1_80}a) and 60\%, 100\% and 120\% (Figure \ref{fig:Run1_80}b-d respectively) for Run 1 and 2. The color scale for Figure \ref{fig:Run1_80}b-d represents the percentage of root pixels in each spatial bin. Figure \ref{fig:Run1_80}e shows the EMD score overall. Figure \ref{fig:Run1_80}f, Figure \ref{fig:Run1_80}g, and Figure \ref{fig:Run1_80}h represent the top 20\% value changes between 80\% and the three other soil water levels; red spatial bins are unique to 80\%, green spatial bins are unique to 60\%, 100\% or 120\%, and yellow spatial bins are common to both water levels.}
    \label{fig:Run1_80}
\end{figure*}

The RSD for the 120\% moisture level appeared extremely different from the other moisture levels (Table \ref{tab:CH_run34}, Figure \ref{fig:Run3_120}). The roots did not grow very deep inside the rhizoboxes, but tended to spread out laterally to a greater extent than in the other soil water conditions. Our STARSEED approach captures this distinct difference between the root images, as most changes between the RSDs are shown through the arrows pointing down from the lateral roots of the 120\% moisture level representative RSD. The root system for 120\% also tended to be denser, as evidenced by the amount of high color value bins that indicate a higher percentage of root pixels in Figure \ref{fig:Run3_120}a. 
\begin{figure*}[htb]
    \centering
    \includegraphics[width=0.95\linewidth]{Figures/Figure8.png}
    \caption{RSD, EMD score and largest value changes between 120\% (Figures \ref{fig:Run3_120}a) and 60\%, 80\% and 100\% (Figures \ref{fig:Run3_120}b-d respectively) for Run 3 and 4. The color scale for Figures \ref{fig:Run3_120}b-d represents the percentage of root pixels in each spatial bin. Figures \ref{fig:Run3_120}e shows the EMD score overall. Figures \ref{fig:Run3_120}f, Figures \ref{fig:Run3_120}g and Figures \ref{fig:Run3_120}h represent the the top 20\% value changes between 120\% and the three other soil water levels; red spatial bins are unique to 120\%, green spatial bins are unique to 60\%, 80\% or 100\%, and yellow spatial bins are common to both water levels.}
    \label{fig:Run3_120}
\end{figure*}

\section*{Discussion}
\subsection*{\textbf{Avoidance vs Tolerance in waterlogged soil conditions}}
One interesting observation is that the three features that were considered (\% root pixels, fractal dimension and lacunarity) had very similar discriminating power between cultivars and between moisture levels (Tables \ref{tab:CH_run12} and \ref{tab:CH_run34}). Both fractal dimension and lacunarity have been used successfully in RSD and root architecture analysis \cite{li2020fractal,walk2004modelling}. However, the main differences between these previous studies and the current one is that in this work these features were not calculated across the entire image but for each spatial bin. By using these texture features to describe the individual regions of an image, we improved the RSD characterization and comparisons for each treatment. While we calculated the fractal dimension and lacunarity values for the whole images, we observed limited differences among treatments for the CH index (\textit{e.g.}, S32 $= 0.06$ and S38 $= 0.03$ for Runs 3 and 4 with fractal dimension). The Kruskal-Wallis/Dunnett’s test with Bonferroni’s adjustment showed that \% root pixels appears to be a better feature than fractal dimension and lacunarity, as \% root pixels led to the highest CH score for images from Runs 3 and 4 across treatments. For Runs 1 and 2, fractal dimension was the highest ranked feature for cultivar, while \% root pixels was better for water levels. Therefore, for future similar work, \% root pixels, an easily calculated parameter, may be preferential to fractal dimension and lacunarity for distinguishing between sesame RSDs.

When looking at these results more specifically, we see that without fertilizer, cultivar appears to be the dominant driver of RSD, and that soil water only plays a minimal role (Table \ref{tab:CH_run12} and \ref{tab:CH_run34}). This can be seen in the highest CH index score on average across all cultivars compared with the averages for water level (Table \ref{tab:CH_run12}). S32 was the cultivar with the most distinct RSD, and in Figure \ref{fig:Run1S32}, we are able to hypothesize what made this RSD different from the others. Specifically, the arrows representing the 20\% highest value changes show that the main differences between the S32 and S35 RSDs were located mostly in the lower (deeper) section of the root system (Fig. \ref{fig:Run1S32}f), and that on the contrary the main differences between the S32 and S40 RSDs were situated in the top (shallower) section of their root systems.

The situation is reversed when fertilizer is added, and soil water level becomes the factor with the biggest impact on RSD, as evidenced by the much higher CH index value for the soil water level treatment. This high score is actually driven by a single treatment, 120\% soil water level, which had a very different RSD compared to the other soil water treatment (Figure \ref{fig:Run3_120}). The roots appeared to not grow as deep and had an increased lateral growth compared to the RSD for the three other water levels. This can be seen through the location of the white arrows, all pointing at changes between the RSD at 120\% and the three other RSDs. This very distinct RSD in flooded conditions when fertility is adequate in Runs 3 and 4 contrasts strongly with that of Run 1 and 2. 
Sesame is known to be highly sensitive to waterlogged soil \cite{sarkar2016effect}, though this sensitivity does not always translate into early total root length differences \cite{gloaguen2021plasticity}. Instead, we see here that the plant seems to be able to activate a variety of morpho-anatomical responses to cope with flooding stress, echoing the literature \cite{wei2013morpho}. Here, we can highlight two different morpho-anatomical strategies employed by the crop depending on the soil fertility status. Without adequate nutrient availability, the crop seems to opt for a \textit{tolerance} strategy to flooding, developing roots into the bottom part of the rhizobox where the soil was saturated with water (Figure \ref{fig:Run1_80}). We can assume that this apparent tolerance strategy is accompanied but other changes not captured in this study such as the formation of aerenchyma and changes in the enzymatic activities \cite{wei2013morpho}. However, when there are enough nutrients in the soil, the plants seemed to adopt an \textit{avoidance} strategy, and did not grow roots down into the waterlogged soil, but tended to proliferate laterally (Figure \ref{fig:Run3_120}). We note that these observations are constant across the cultivars; as a result, fertility may condition the response of early sesame RSDs to flooding stress. 

\subsection*{\textbf{Validity of the EMD Method}}
Although of interest in estimating early root vigor and biomass, only considering TRL provides limited insight about RSD as TRL does not consider the spatial arrangements of the roots \cite{gloaguen2021plasticity}. Increasing our knowledge and understanding of root development in response to environmental stresses is of capital importance for global agricultural production, and has been defined as the pillar of the Second Green Revolution \cite{lynch2007roots}. Many methods are now available to observe RSD and root architecture, yet some of the more advanced techniques such as CT imaging remain expensive and impractical, underlining the need for improved trans-disciplinary phenotyping approaches that can capture and quantify the complexity of RSD and root system architecture \cite{atkinson2019uncovering,wu2018optimizing,zhu2011lab}. 

Our proposed STARSEED provides a visual, explicit, and quantitative characterization of RSD, allowing for spatially precise comparisons between RSD. A valuable aspect of the analysis is the introduction of the EMD class score, which summarizes all the differences between two RSDs down to a single number. The EMD method, combined with the CH index, can be used to very quickly know which treatment lead to the most distinct RSD, and the degree to which these RSDs relate to one another globally. The STARSEED method could also be used in combination with other RSD and root system architecture measurements such as branching, angle, hierarchy, and fine-root distribution to achieve a comprehensive mechanistic understanding of root system.

The actual spatial EMD result (\textit{i.e.}, Figures \ref{fig:Run1S32} through \ref{fig:Run3_120}) can then used to further elucidate specific differences between RSDs. To the best of the authors' knowledge, this is the first time EMD has been used to characterize RSD. The method has been thoroughly validated and used in varied fields of science, including fluid mechanics \cite{benamou2000computational}, linguistics \cite{kusner2015word}, or, more related to the present study, image classification and comparison \cite{zhang2020deepemd}. The CH index used in STARSEED serves as an evaluation step for the method to not only quantify what we observe, but also to ensure the approach captures distinct features to effectively group RSD based on shared spatial characteristics, which can then be linked to the treatment structure of the experiment. 

One of the most critical strengths of STARSEED is that the proposed method allows for the generation of an ``average" RSD visual representation called the representative RSD image for that treatment, as shown in Figures \ref{fig:Run1S32} through \ref{fig:Run3_120}a through d. By partitioning this newly generated image into local bins and calculating a feature value for each of these spatial bins, we can generate a visually explicit average RSD that can be compared to other representatives RSD to study differences between genotypes or environmental conditions (or any other treatment given a different experiment). Recent studies have attempted to either develop new techniques to directly measure RSD and/or root system architecture \cite{armengaud2009ez}, find a way to accurately represent average root systems corresponding to specific growing conditions \cite{shahzad2018ez}, and many more studies have developed or refined root development models \cite{tron2015can,postma2017opensimroot,zhao2017root,pages2020metamodelling}. One study in particular created similar 2D heat maps of ``root frequency" observed on the four transparent surfaces of rhizoboxes but did not perform a spatially explicit analysis to the level of STARSEED \cite{jorgensen2014spatial}. Furthermore, the STARSEED approach is able to characterize and compare 2D RSD images of root systems at various scales. RSD is typically measured through soil coring at various depths, using a grid on a profile wall, or with mini-rhizotron systems \cite{zhi2017root,LI201747,li2017effects}. These techniques lack an efficient way to integrate local data at the whole root system's scale across all depths. STARSEED provides a way to get information about the entire RSD (with the outputs presented in Figures \ref{fig:Run1S32} through \ref{fig:Run3_120} a, b, c and d) and at the same time highlight the largest RSD differences between two RSD signatures (Figures \ref{fig:Run1S32} through \ref{fig:Run3_120} f, g, and h). This feature is quite unique and could be of great value to breeding programs where RSD need to be compared. 
Some of the shortcomings of STARSEED include the need to hand-label root images with WinRhizoTron. This is an extremely time consuming task that represents a serious limit to the rapidity and ease with which our approach can be used. However, recent work in machine learning is paving the way for fully automated and robust root tracing algorithms \cite{yu2020weakly}, which would dramatically increase the applicability of STARSEED. In addition, STARSEED remains an indirect measure of similarity between root systems as the proposed method is based solely on image analysis. STARSEED only indicates relative similarity or differences between two root systems captured with the same imaging system.

\subsection*{\textbf{Going beyond the 2D images of rhizoboxes}}
Given the rhizobox set up of this study and the use of flatbed scanners for root images acquisition, the observations are confined to 2D when root systems always exist in 3D spaces. Imaging and characterizing a whole root system in 3D \textit{in situ} or even \textit{in vitro} is usually low throughput and expensive, if not nearly impossible. Most scientists thus resort to using models in lieu of direct observations, as previously mentioned. One of the main difficulty of using these 3D root development models is correct parameterization, which is crucial to the accuracy and validity of a model's prediction \cite{schnepf2018statistical}. Recent work has shown that 2D measurements of root systems could be used to adequately inform the parameters for 3D models in wheat \cite{landl2018measuring}. It is thus a reasonable hypothesis to say that the current STARSEED approach could be further refined and subsequently used to generate and interpret such 3D models. There is potential to go even further and directly use STARSEED, and more broadly EMD, on 3D data \cite{salti2014shot}, although acquiring such data still remains challenging. 

Combining the average EMD visual maps with new RSD and root system architecture characterization techniques \cite{wu2014integrated} have the potential to boost our understanding of root system plasticity. These tools would allow for both global (\textit{i.e.}, EMD scores) and local (\textit{i.e.}, magnitude and direction of changes) comparisons between genotypes and environmental conditions. {We can also quantify the distribution of arrows as this information is captured from the flow matrix. We could record the distribution of the magnitude and direction of the arrows (similar to histogram-based features such as histogram of oriented gradients \mbox{\cite{dalal2005histograms}} or edge histogram descriptors \mbox{\cite{frigui2008detection}}) for further analysis.} Another interesting direction would be to develop a way to connect the observations made with STARSEED to more mechanistic measurements.


\section*{Conclusion}\label{sect:conclusion}
In this paper, we presented STARSEED, an approach to characterize root system distribution from 2D images. Qualitative and quantitative analysis demonstrate the effectiveness of the proposed method. STARSEED successfully incorporated spatial and texture information to describe root architectures in both global and local contexts. The method is explainable and provides clear connections to biological aspects of each root image. STARSEED also allows for aggregating individual architectures to assess and average responses for each environmental condition and genotype. Future work includes clustering based on root architecture (\textit{i.e.}, using the pairwise EMD matrix for relational clustering), {comparative study of different measures (\textit{e.g.}, convex hull) for feature extraction}, applying our general framework over time to characterize how RSA develops in various conditions, along with scaling up to 3D characterization.


\section*{Appendix}

%%%%%%%%%%%%%%%%%%%%%%%%%%%%%%%%%%%%%%%%%%%%%%
%%                                          %%
%% Backmatter begins here                   %%
%%                                          %%
%%%%%%%%%%%%%%%%%%%%%%%%%%%%%%%%%%%%%%%%%%%%%%

\begin{backmatter}

\section*{Acknowledgements}%% if any

\section*{Funding}%% if any
This material is based upon work supported by the National Science Foundation Graduate Research Fellowship under Grant No. DGE-1842473.

\section*{Abbreviations}%% if any
Not applicable 

\section*{Availability of data and materials}%% if any
The raw and binarized root images for our experiments are publicly available: \url{https://github.com/GatorSense/STARSEED/tree/master/Data}.

\section*{Ethics approval and consent to participate}%% if any
Not applicable 

\section*{Competing interests}
The authors have no conflicts of interest 
to declare that are relevant to the content of this article.

\section*{Consent for publication}%% if any
Not applicable

\section*{Authors' contributions}
Joshua Peeples, Weihuang Xu, Romain Gloaguen, Diane Rowland, and Alina Zare planned and designed the research. Joshua Peeples, Weihuang Xu, and Romain Gloaguen collected data, conducted experiments, and performed analysis. Joshua Peeples, Weihuang Xu, and Romain Gloagen wrote the manuscript. Diane Rowland, Alina Zare, and Zachary Brym reviewed and edited the manuscript. All authors read and approved the final manuscript.

% \section*{Authors' information}%% if any
% Text for this section\ldots

%%%%%%%%%%%%%%%%%%%%%%%%%%%%%%%%%%%%%%%%%%%%%%%%%%%%%%%%%%%%%
%%                  The Bibliography                       %%
%%                                                         %%
%%  Bmc_mathpys.bst  will be used to                       %%
%%  create a .BBL file for submission.                     %%
%%  After submission of the .TEX file,                     %%
%%  you will be prompted to submit your .BBL file.         %%
%%                                                         %%
%%                                                         %%
%%  Note that the displayed Bibliography will not          %%
%%  necessarily be rendered by Latex exactly as specified  %%
%%  in the online Instructions for Authors.                %%
%%                                                         %%
%%%%%%%%%%%%%%%%%%%%%%%%%%%%%%%%%%%%%%%%%%%%%%%%%%%%%%%%%%%%%

% if your bibliography is in bibtex format, use those commands:
\bibliographystyle{bmc-mathphys} % Style BST file (bmc-mathphys, vancouver, spbasic).
\bibliography{bmc_article}      % Bibliography file (usually '*.bib' )
% for author-year bibliography (bmc-mathphys or spbasic)
% a) write to bib file (bmc-mathphys only)
% @settings{label, options="nameyear"}
% b) uncomment next line
%\nocite{label}

% or include bibliography directly:
% \begin{thebibliography}
% \bibitem{b1}
% \end{thebibliography}

%%%%%%%%%%%%%%%%%%%%%%%%%%%%%%%%%%%
%%                               %%
%% Figures                       %%
%%                               %%
%% NB: this is for captions and  %%
%% Titles. All graphics must be  %%
%% submitted separately and NOT  %%
%% included in the Tex document  %%
%%                               %%
%%%%%%%%%%%%%%%%%%%%%%%%%%%%%%%%%%%

% %%
% %% Do not use \listoffigures as most will included as separate files

\newpage
\section*{Supplemental Figures}
\renewcommand{\thefigure}{S\arabic{figure}}
\setcounter{figure}{0}
  \begin{figure}[h!]
    \centering
    \includegraphics[width=0.90\linewidth]{Figures/FigureS1.png}
  \caption{Average computational cost for Runs 1 and 2. The shaded areas correspond to $\pm$ 1 standard deviation across three experimental runs of each feature and grid size value. The experiments were performed on a Dell XPS 15 9520 laptop with 12th Generation Intel Core i9-12900 HK processor.
}
\label{fig:S1}
\end{figure}

  \begin{figure}[h!]
    \centering
    \includegraphics[width=0.90\linewidth]{Figures/FigureS2.png}
  \caption{Average computational cost for Runs 3 and 4. The shaded areas correspond to $\pm$ 1 standard deviation across three experimental runs of each feature and grid size value. The experiments were performed on a Dell XPS 15 9520 laptop with 12th Generation Intel Core i9-12900 HK processor.
}
\label{fig:S2}
\end{figure}


\begin{figure*}[th]
     \centering
     \begin{subfigure}[b]{0.22\textwidth}
         \centering
         \includegraphics[width=\textwidth]{Figures/FigureS3a.jpg}
         \caption{Cultivar S32}
     \end{subfigure}
     \hfill
     \begin{subfigure}[b]{0.22\textwidth}
         \centering
         \includegraphics[width=\textwidth]{Figures/FigureS3b.jpg}
         \caption{Cultivar S35}
     \end{subfigure}
     \hfill
     \begin{subfigure}[b]{0.22\textwidth}
         \centering
         \includegraphics[width=\textwidth]{Figures/FigureS3c.jpg}
         \caption{Cultivar S38}
     \end{subfigure}
      \hfill
     \begin{subfigure}[b]{0.22\textwidth}
         \centering
         \includegraphics[width=\textwidth]{Figures/FigureS3d.jpg}
         \caption{Cultivar S40}
     \end{subfigure}
        \caption{RGB root images of different cultivars with the same treatment of $60\%$ water level.}
        \label{fig:S3}
\end{figure*}

\begin{figure*}[th]
     \centering
     \begin{subfigure}[b]{0.21\textwidth}
         \centering
         \includegraphics[width=\textwidth]{Figures/FigureS4a.jpg}
         \caption{Water level $60\%$}
     \end{subfigure}
     \hfill
     \begin{subfigure}[b]{0.21\textwidth}
         \centering
         \includegraphics[width=\textwidth]{Figures/FigureS4b.jpg}
         \caption{Water level $80\%$}
     \end{subfigure}
     \hfill
     \begin{subfigure}[b]{0.21\textwidth}
         \centering
         \includegraphics[width=\textwidth]{Figures/FigureS4c.jpg}
         \caption{Water level $100\%$}
     \end{subfigure}
      \hfill
     \begin{subfigure}[b]{0.21\textwidth}
         \centering
         \includegraphics[width=\textwidth]{Figures/FigureS4d.jpg}
         \caption{Water level $120\%$}
     \end{subfigure}
        \caption{RGB root images of the same cultivars (S38) with different water level treatments.}
        \label{fig:S4}
\end{figure*}
% %%%%%%%%%%%%%%%%%%%%%%%%%%%%%%%%%%%
% %%                               %%
% %% Tables                        %%
% %%                               %%
% %%%%%%%%%%%%%%%%%%%%%%%%%%%%%%%%%%%

% %% Use of \listoftables is discouraged.
% %%
% \section*{Tables}
% \begin{table}[h!]
% \caption{Sample table title. This is where the description of the table should go}
%   \begin{tabular}{cccc}
%     \hline
%     & B1  &B2   & B3\\ \hline
%     A1 & 0.1 & 0.2 & 0.3\\
%     A2 & ... & ..  & .\\
%     A3 & ..  & .   & .\\ \hline
%   \end{tabular}
% \end{table}

% %%%%%%%%%%%%%%%%%%%%%%%%%%%%%%%%%%%
% %%                               %%
% %% Additional Files              %%
% %%                               %%
% %%%%%%%%%%%%%%%%%%%%%%%%%%%%%%%%%%%

% \section*{Additional Files}
%   \subsection*{Additional file 1 --- Sample additional file title}
%     Additional file descriptions text (including details of how to
%     view the file, if it is in a non-standard format or the file extension).  This might
%     refer to a multi-page table or a figure.

%   \subsection*{Additional file 2 --- Sample additional file title}
%     Additional file descriptions text.

\end{backmatter}
\end{document}
