## Supplementary figures and images for "Spatial and Texture Analysis of Root System Distribution with Earth Mover’s Distance (STARSEED)"

### Figure1.png

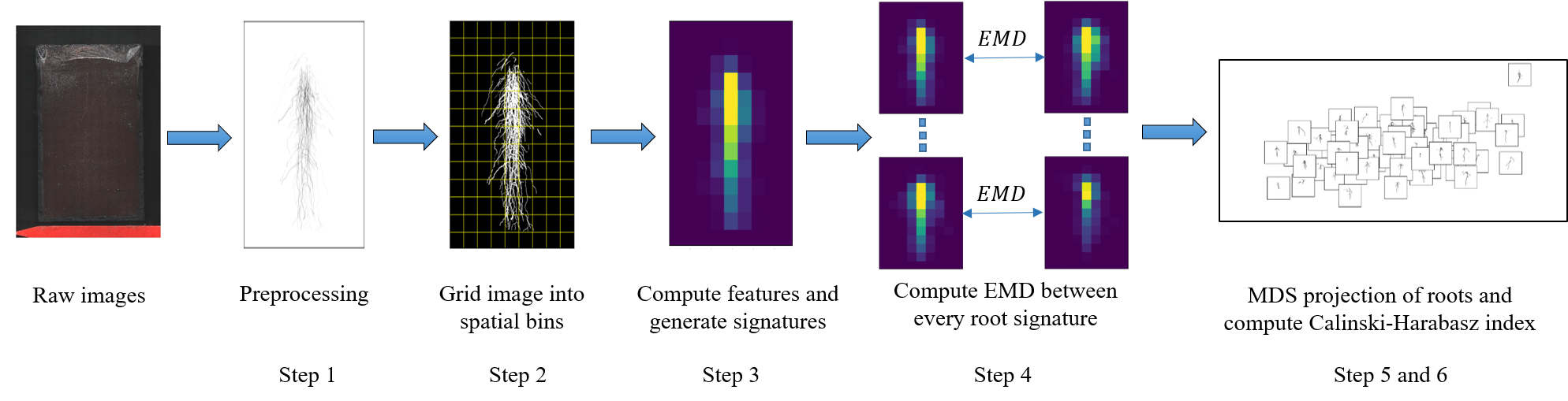

### Figure2.jpg

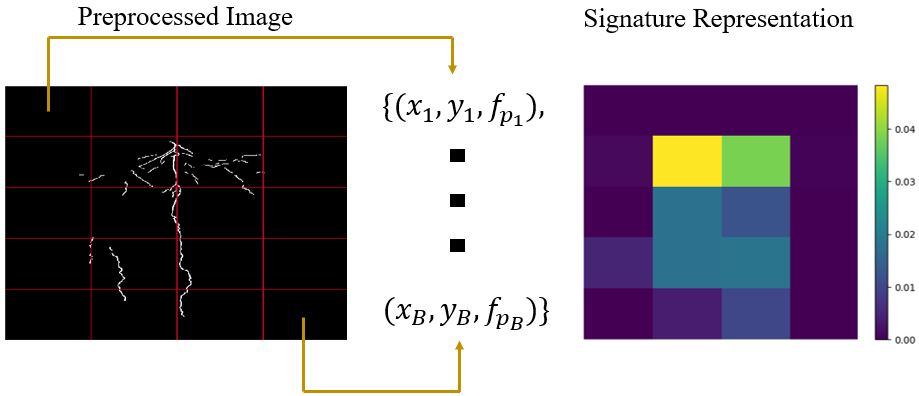

### Figure3.png

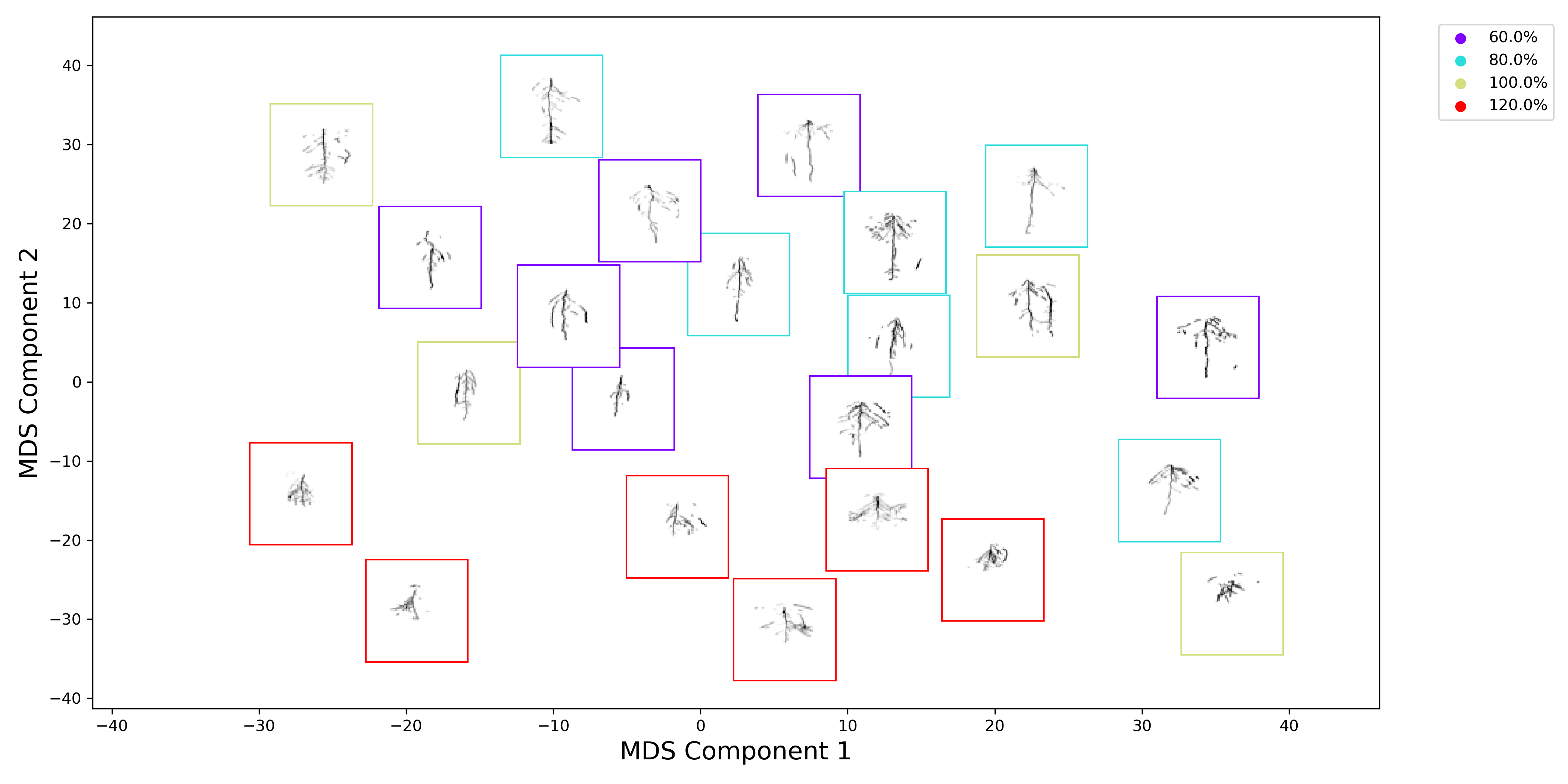

### Figure4.png

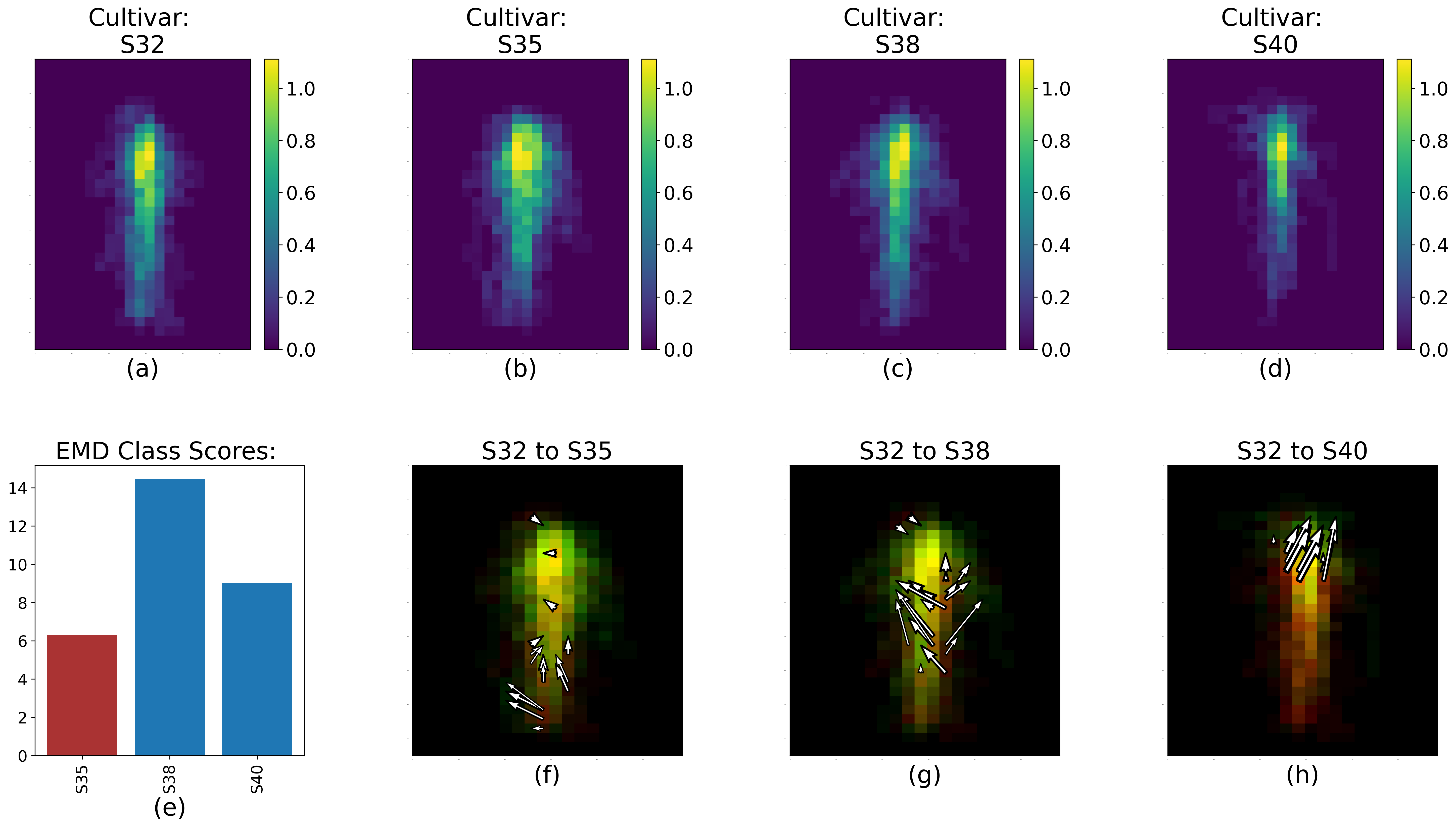

### Figure5.png

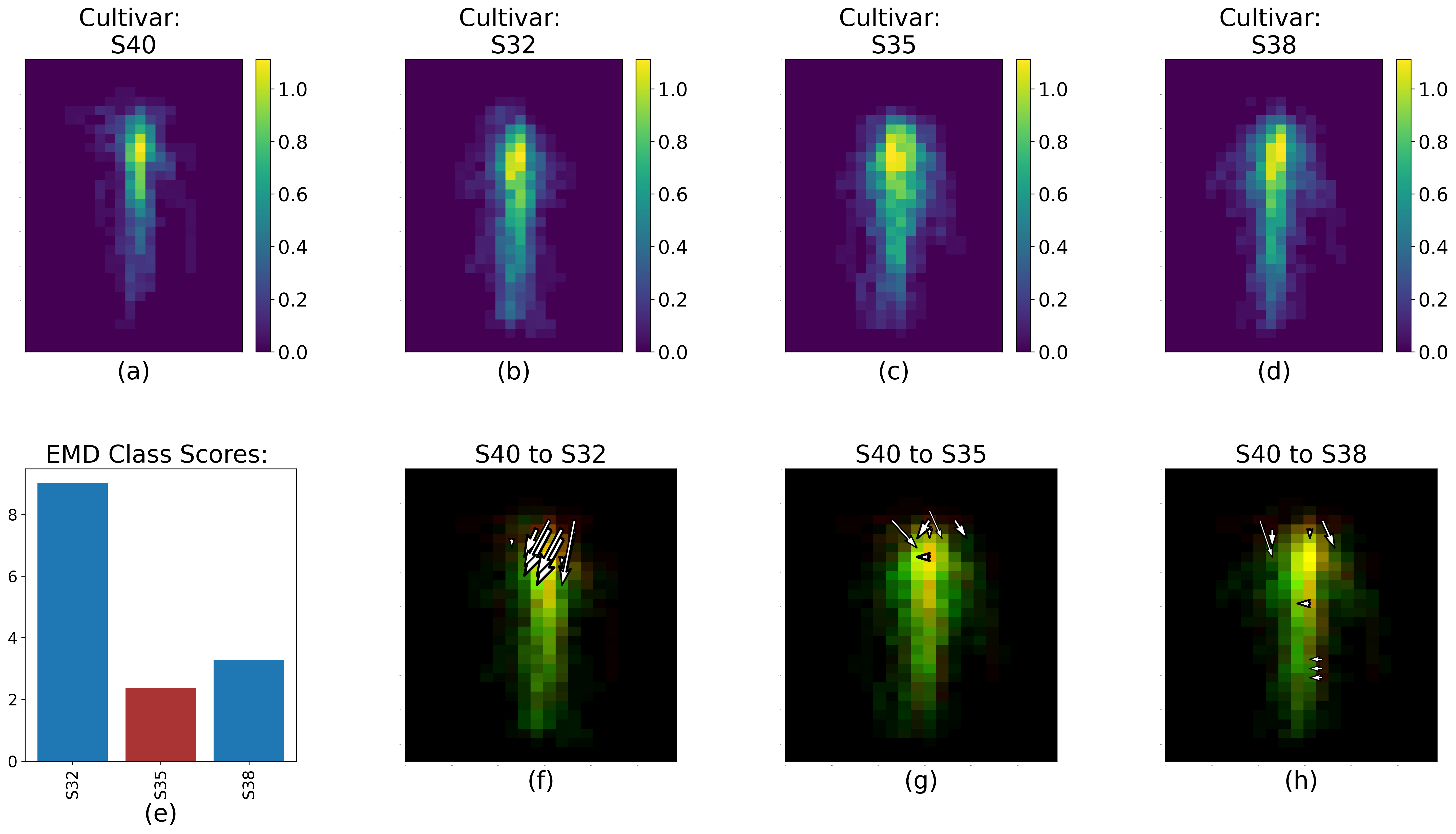

### Figure6.png

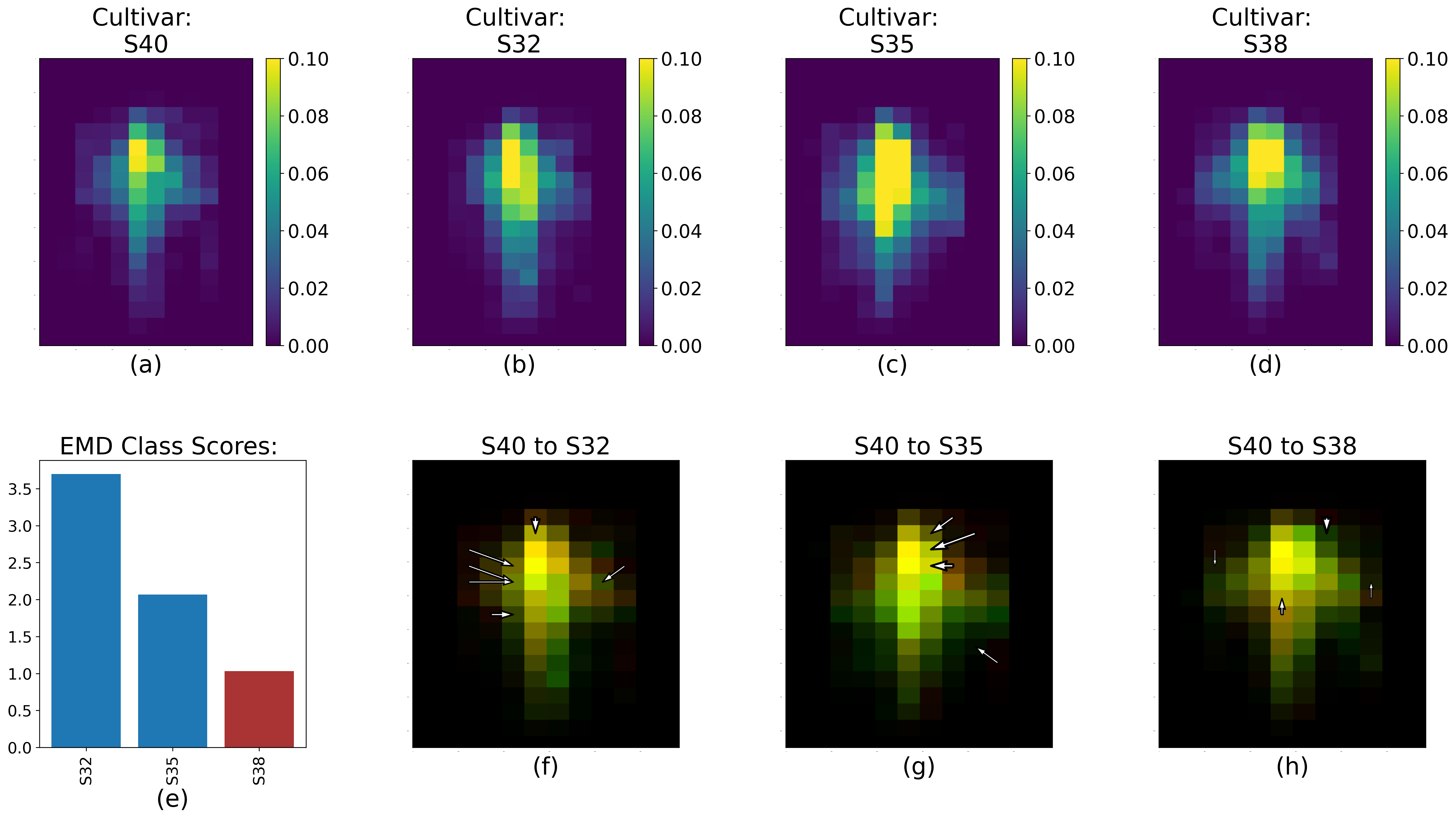

### Figure7.png

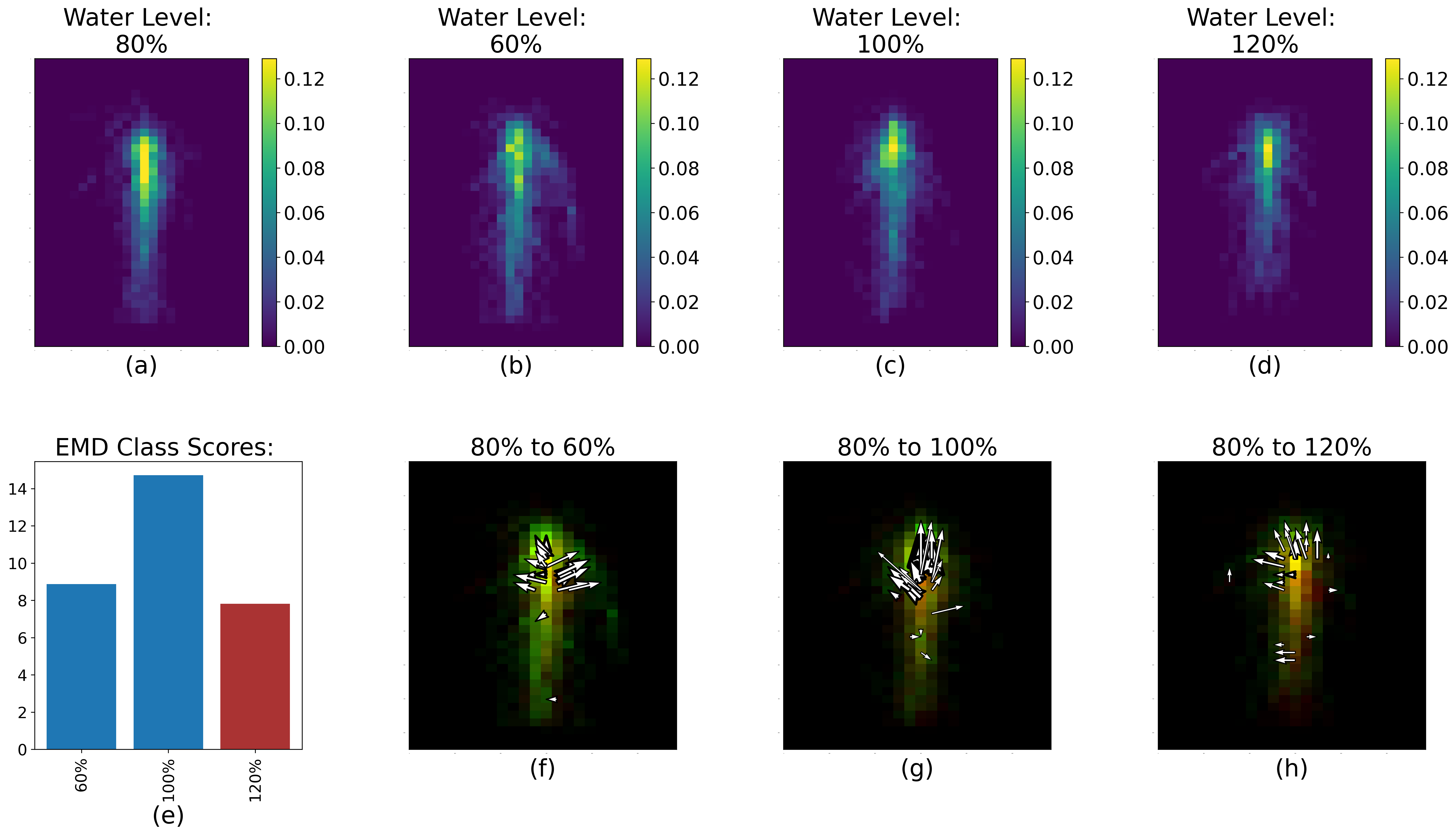

### Figure8.png

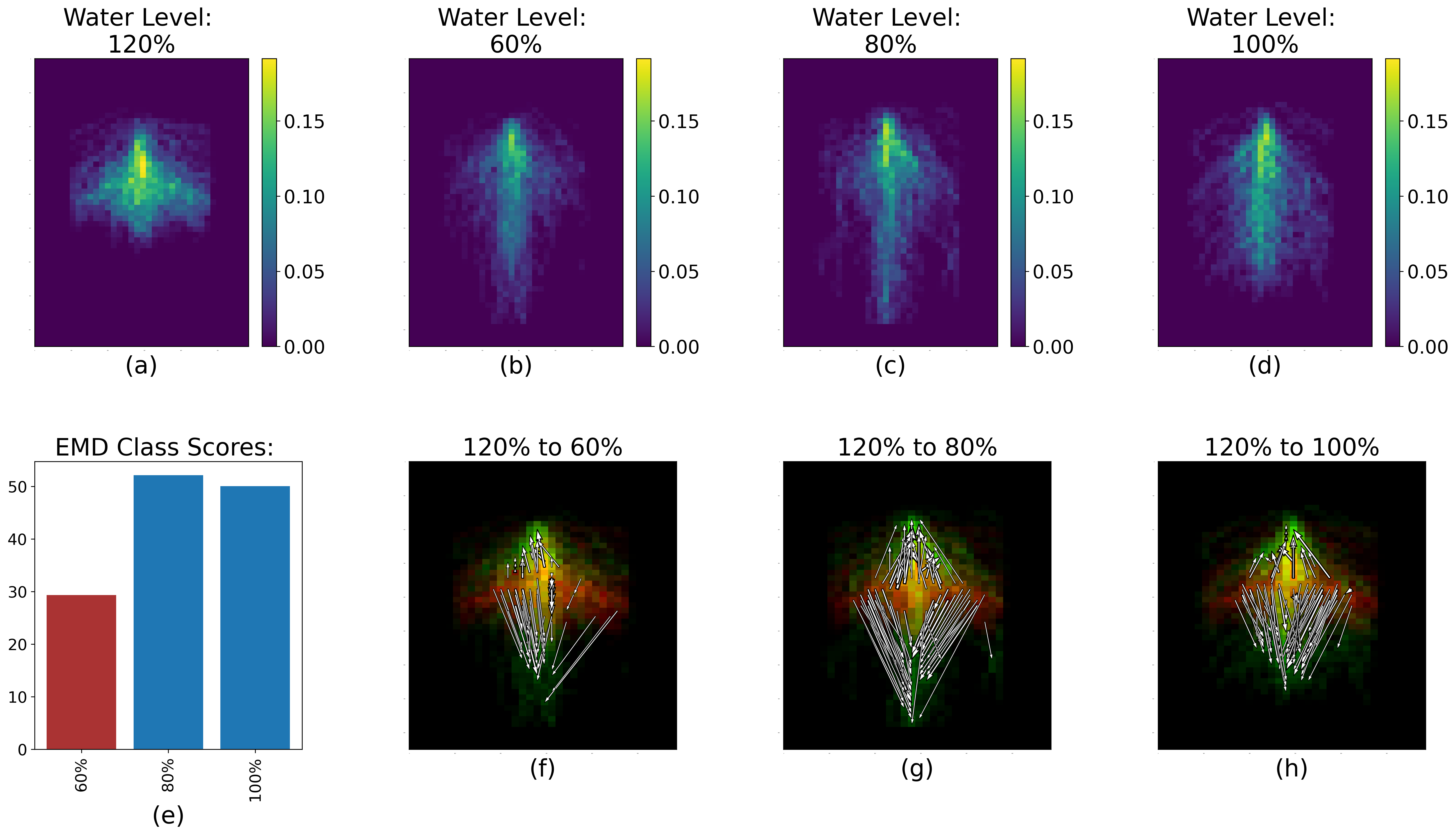

### FigureS1.png

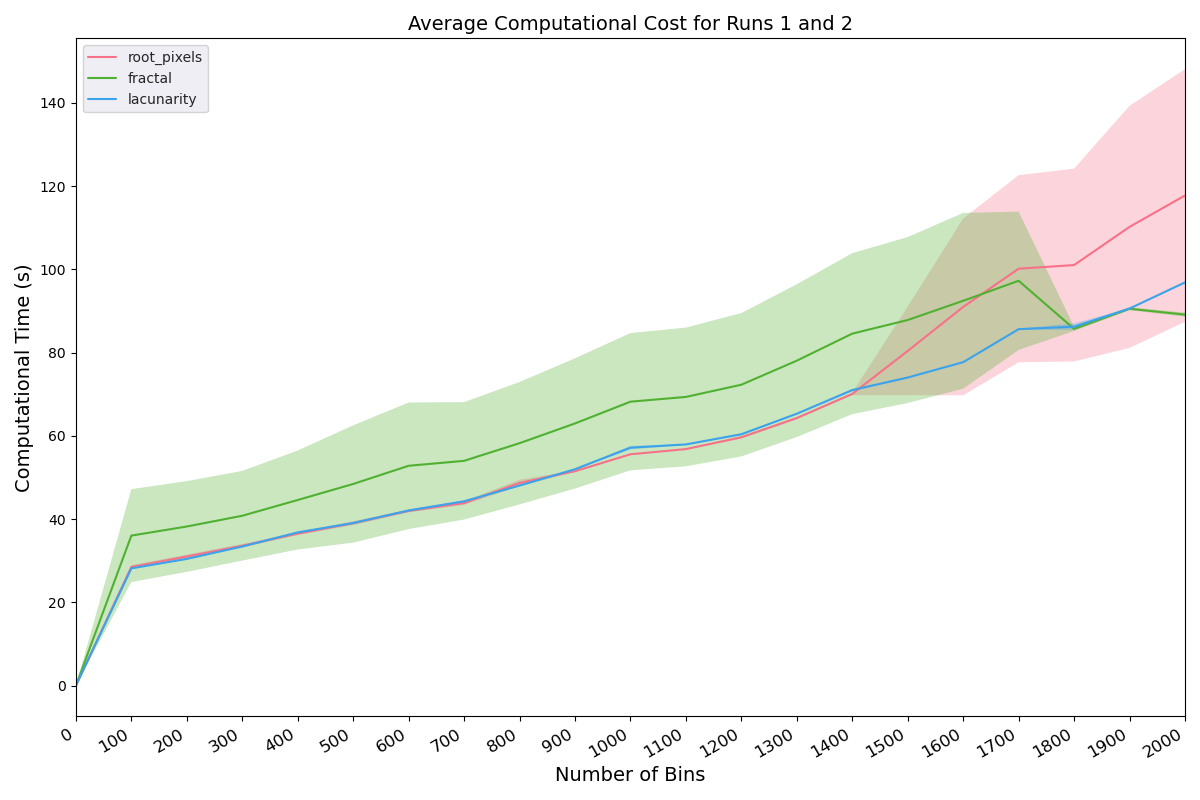

### FigureS2.png

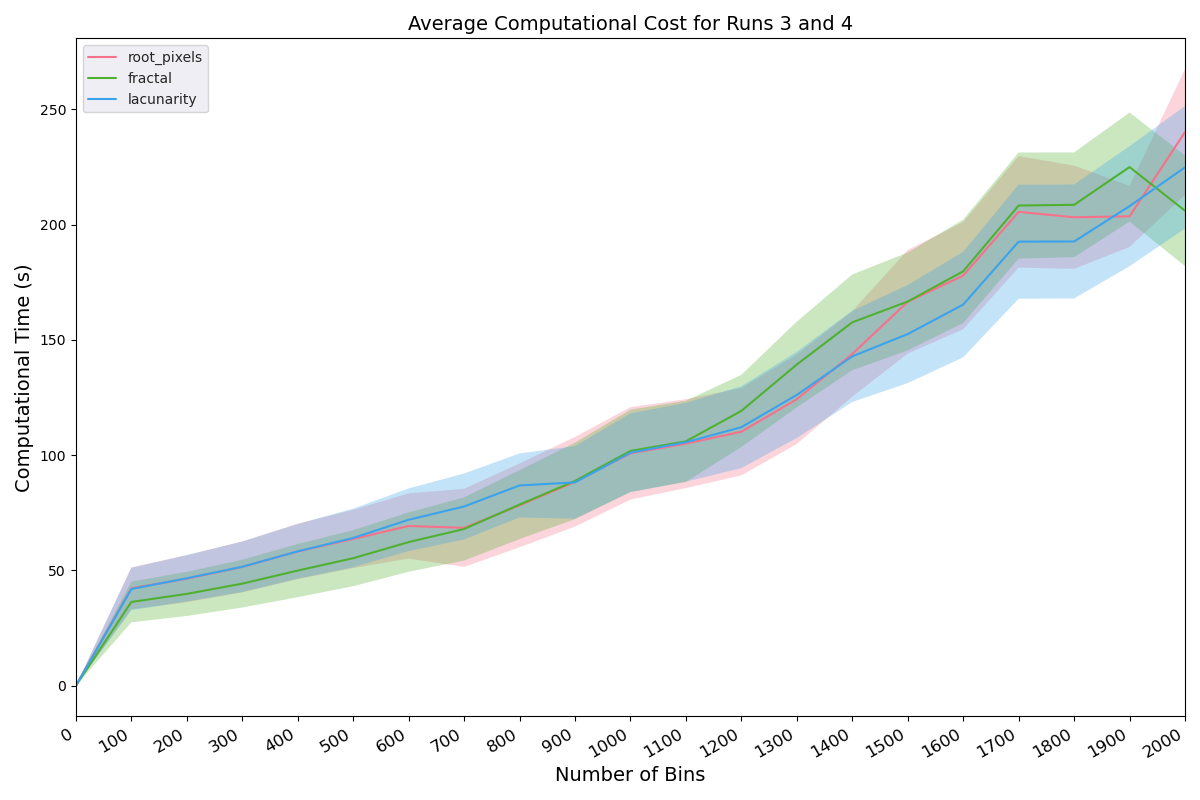

### FigureS3a.jpg

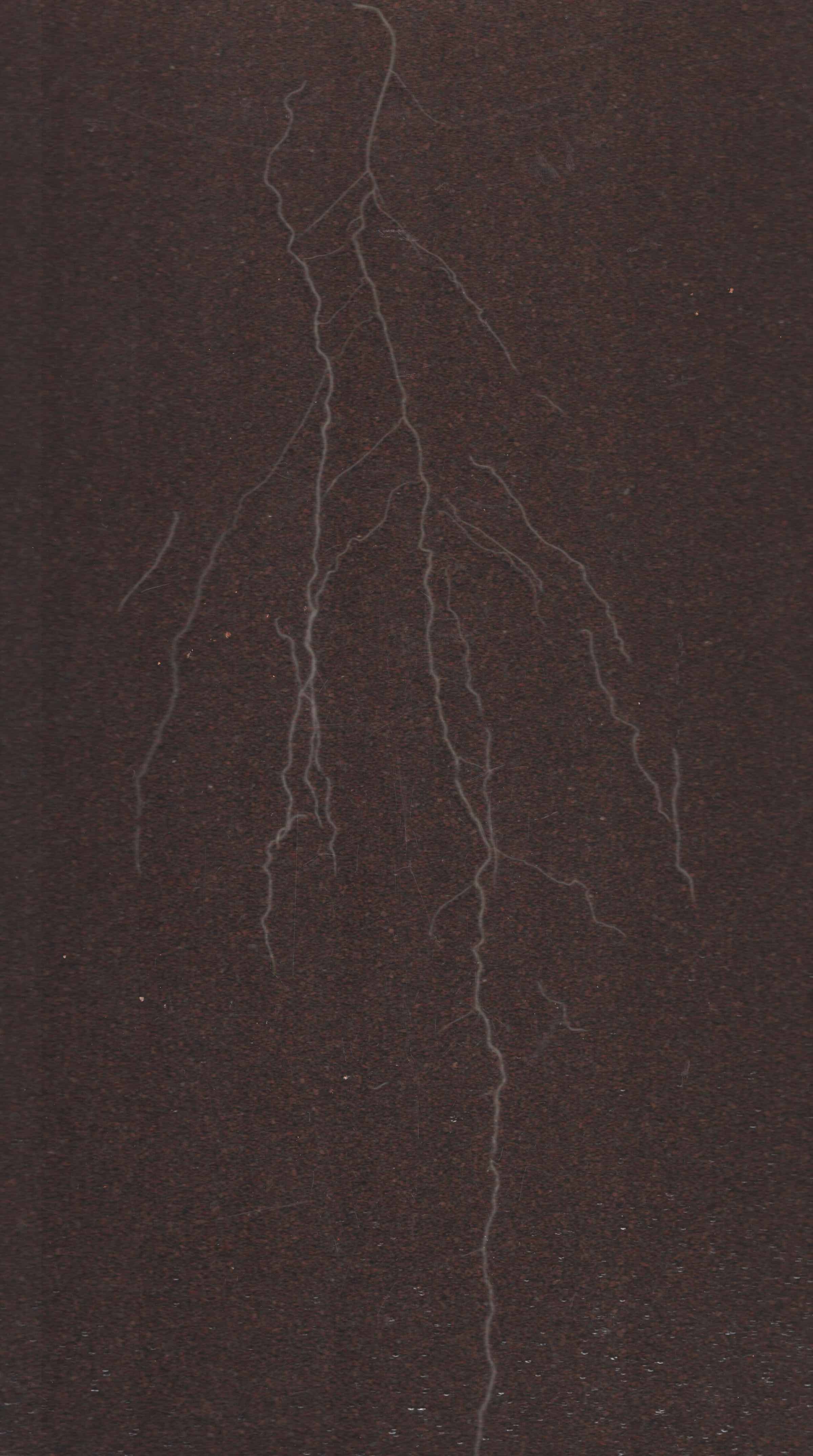

### FigureS3b.jpg

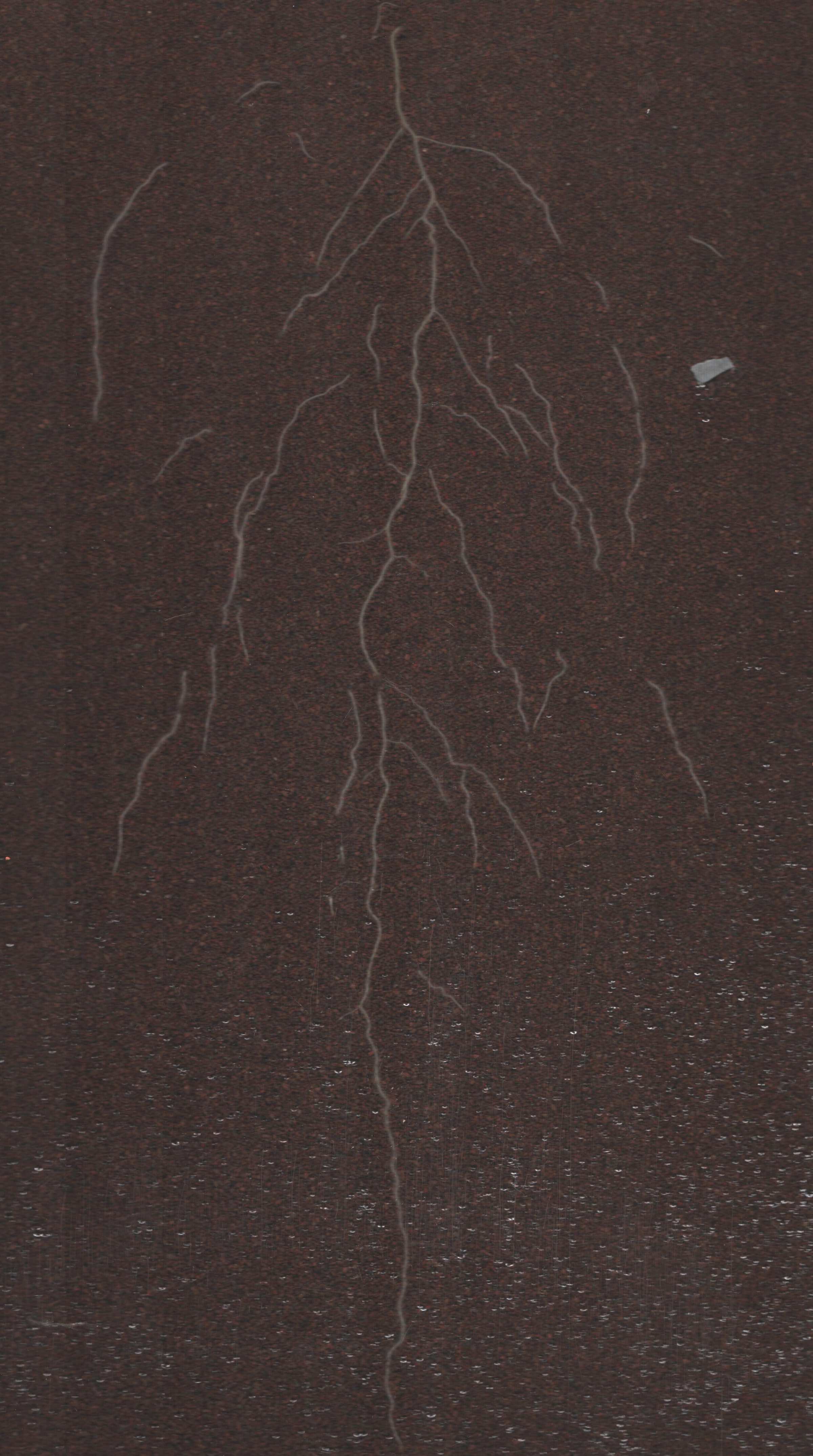

### FigureS3c.jpg

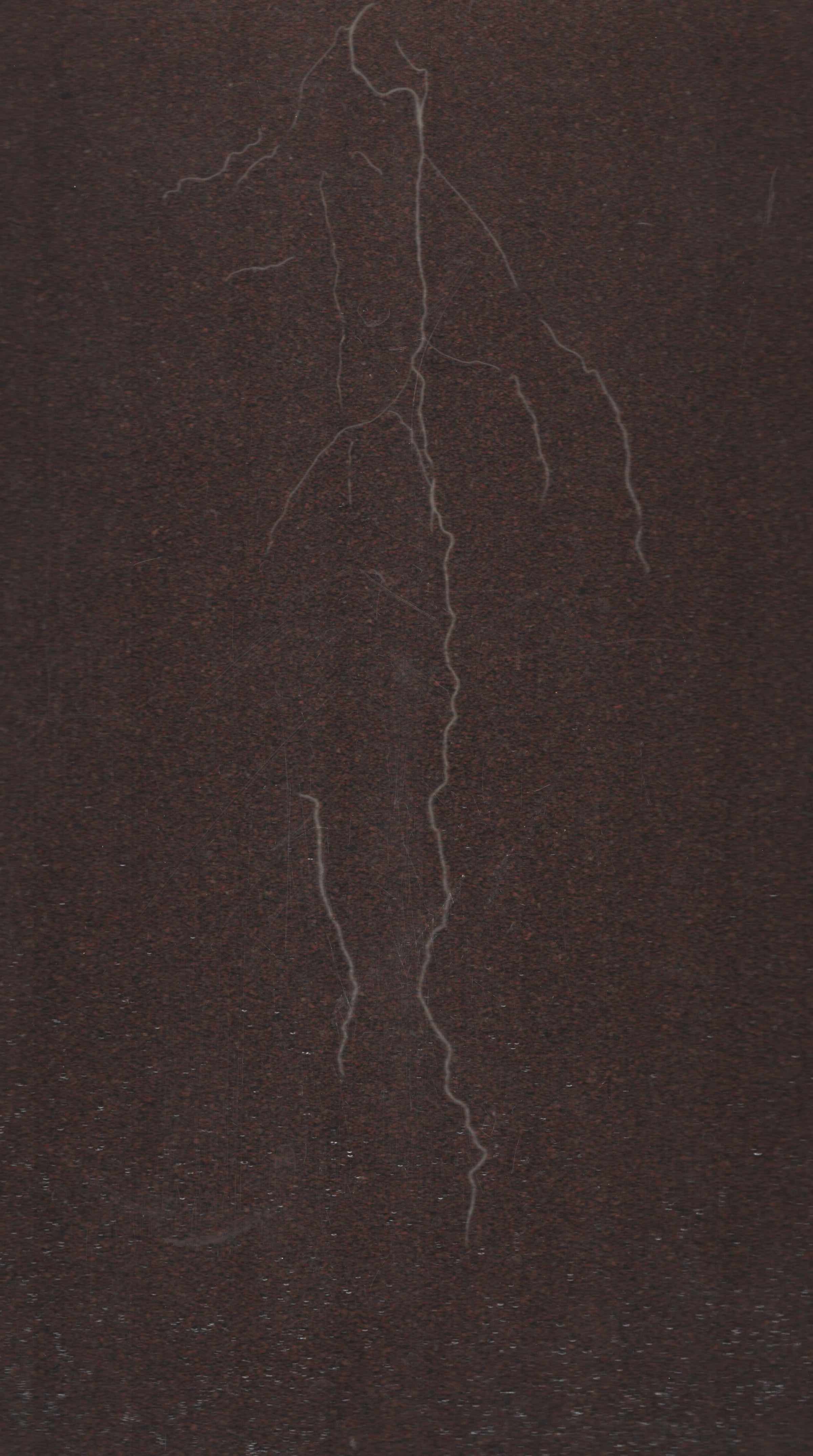

### FigureS3d.jpg

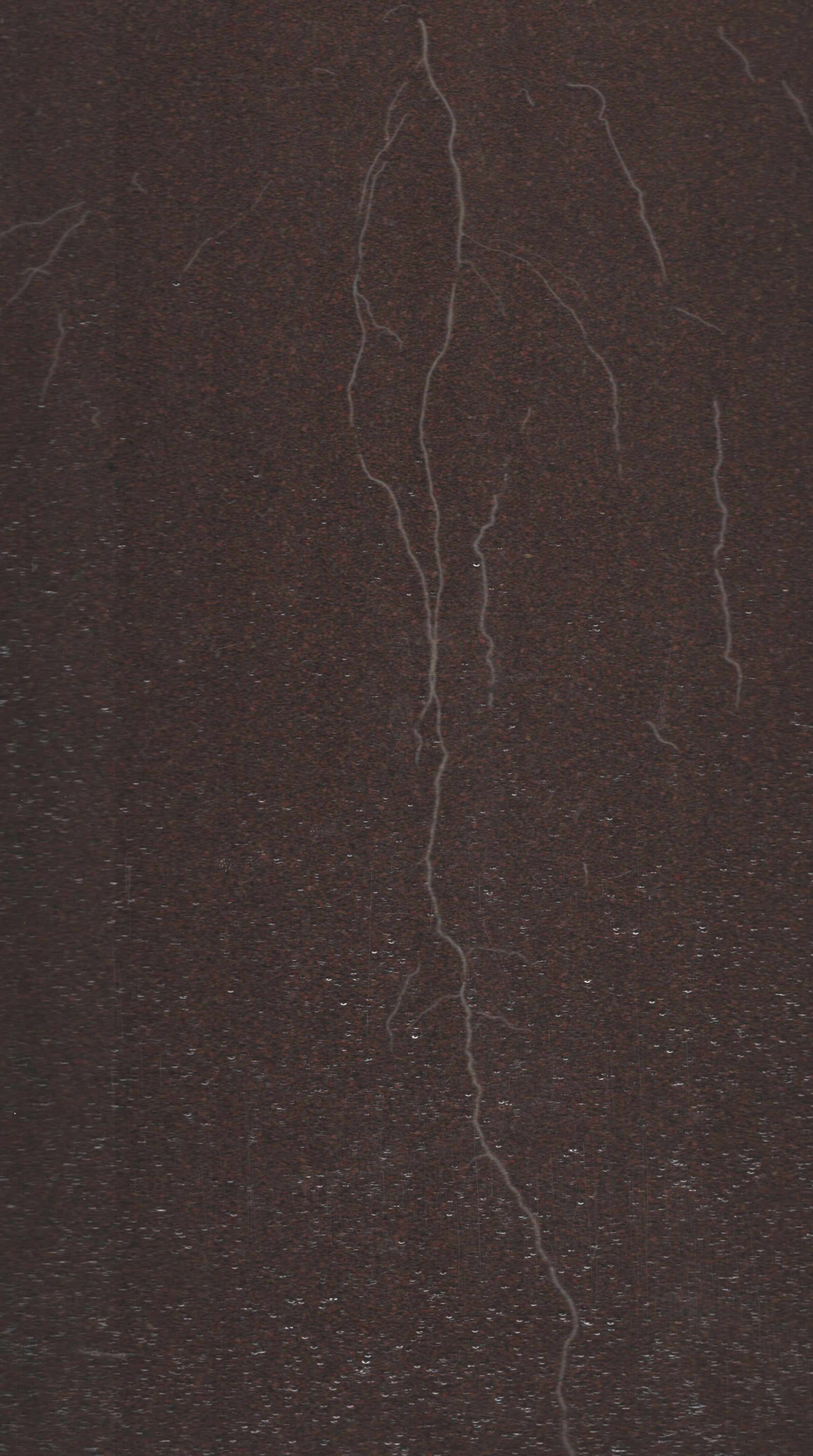

### FigureS4a.jpg

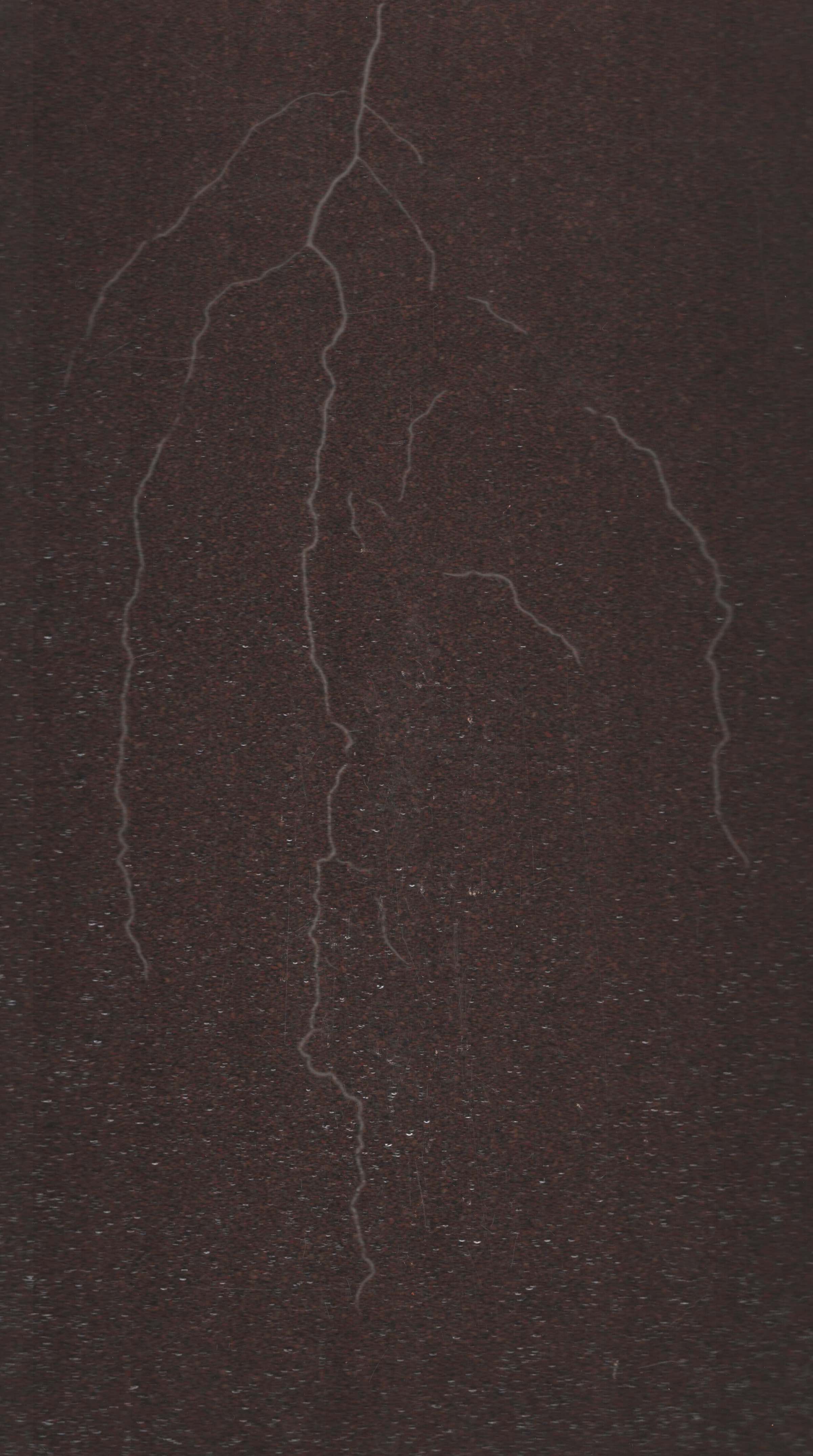

### FigureS4b.jpg

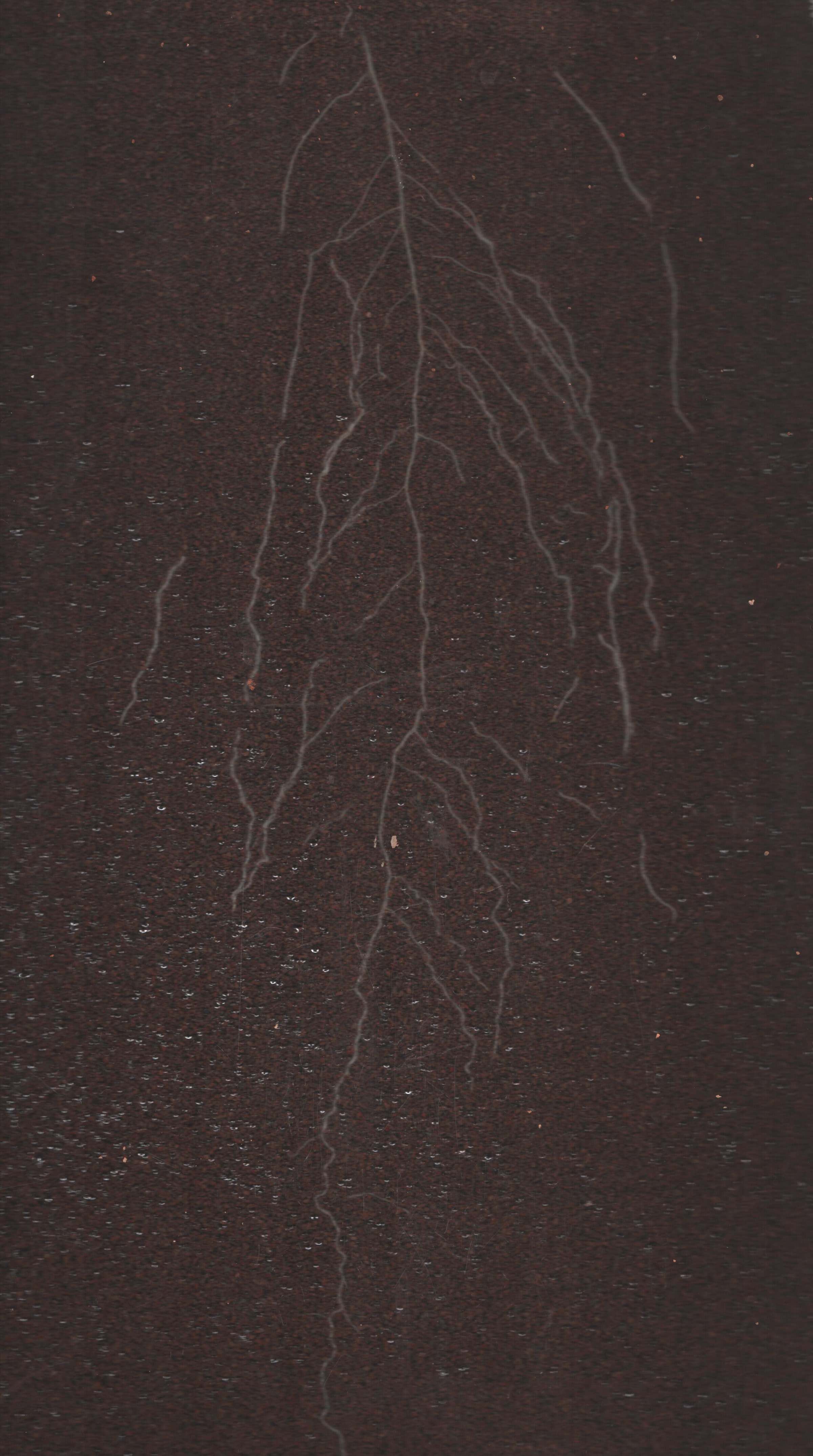

### FigureS4c.jpg

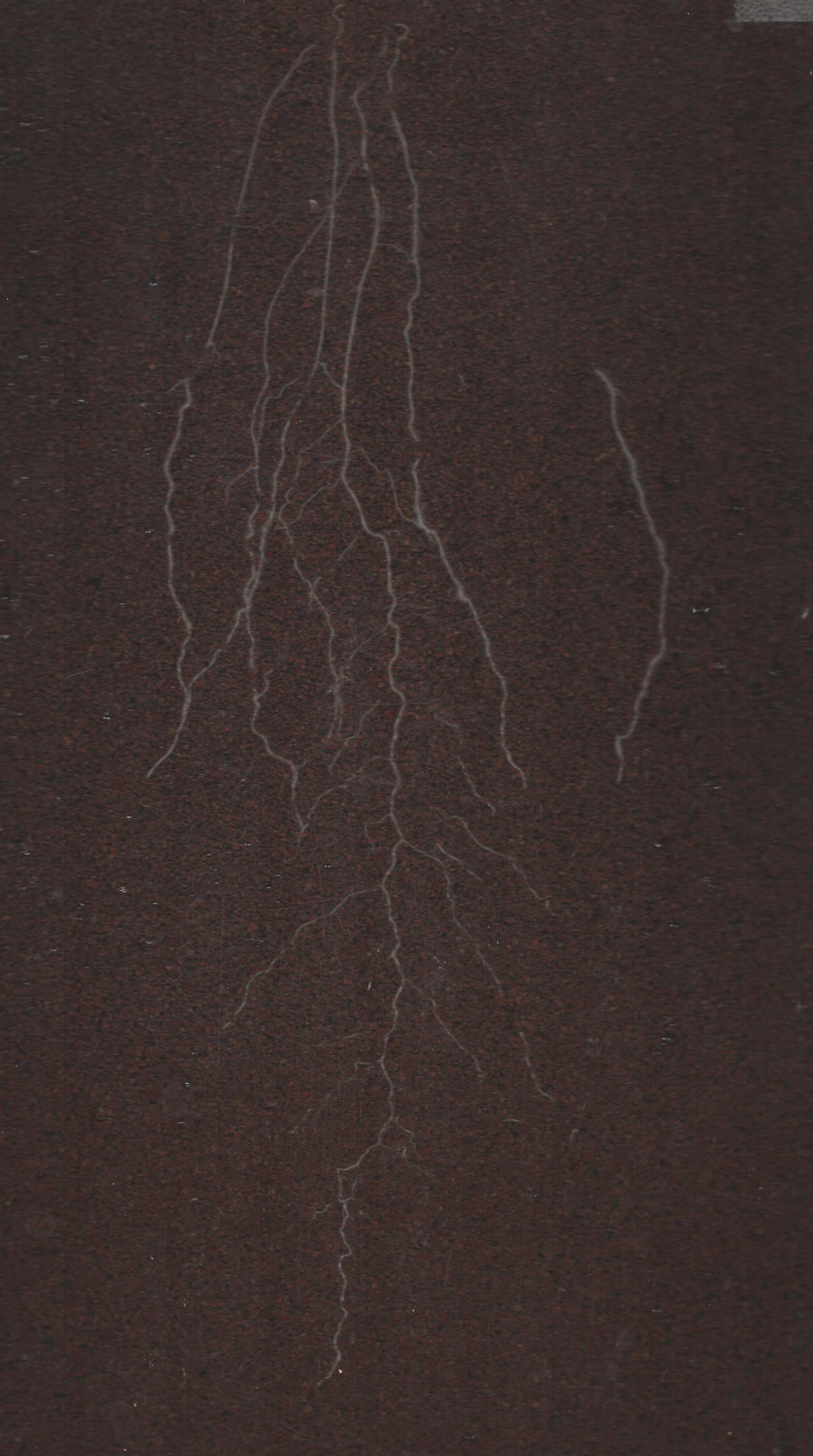

### FigureS4d.jpg

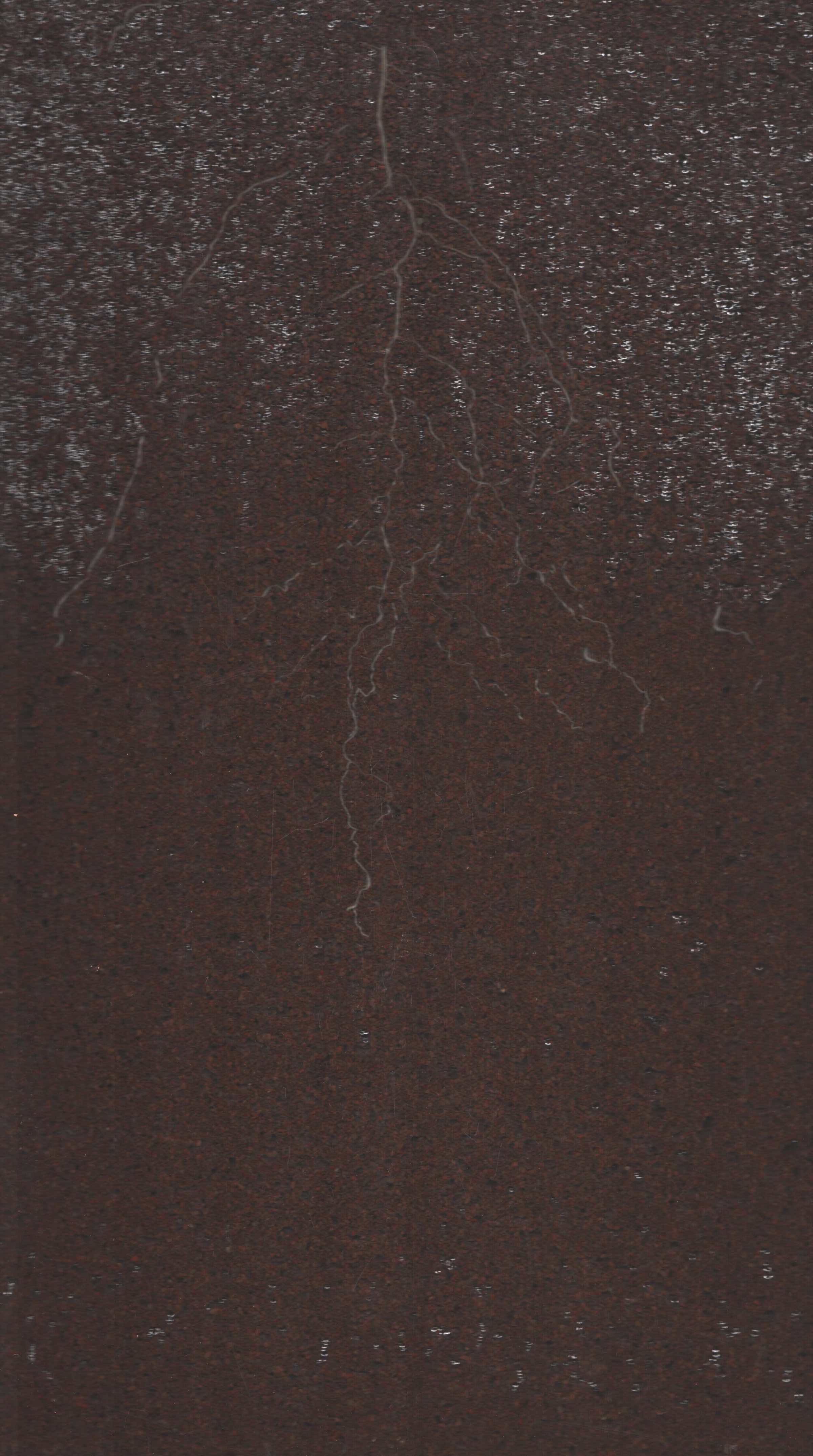
